# Memory for Individual Items is Related to Non-Reinforced Preference Change

**DOI:** 10.1101/621292

**Authors:** Rotem Botvinik-Nezer, Akram Bakkour, Tom Salomon, Daphna Shohamy, Tom Schonberg

## Abstract

It is commonly assumed that memories contribute to value-based decisions. Nevertheless, most theories of value-based decision-making do not account for memory influences on choice. Recently, new interest has emerged in the interactions between these two fundamental processes, mainly using reinforcement-based paradigms. Here, we aimed to study the role memory processes play in preference change following the non-reinforced cue-approach training (CAT) paradigm. In CAT, the mere association of cued items with a speeded motor response influences choices. Previous studies with this paradigm showed that a single training session induces a long-lasting effect of enhanced preferences for high-value trained stimuli, that is maintained for several months. We hypothesized that CAT influences memory accessibility for trained items, leading to enhanced accessibility of their positive associative memories and in turn to preference changes. In two pre-registered experiments, we tested whether memory for trained items was enhanced following CAT, in the short and in the long-term, and whether memory modifications were related to choices. We found that memory was enhanced for trained items and that better memory was correlated with enhanced preferences at the individual item level, both immediately and one month following CAT. Our findings show that memory plays a central role in value-based decision-making following CAT, even in the absence of external reinforcements. These findings contribute to new theories relating memory and value-based decision-making and set the groundwork for the implementation of novel behavioral interventions that lead to long-lasting behavioral change.

## Introduction

Value-based decision-making and memory are both extensively studied processes in cognitive psychology and cognitive neuroscience (Fellows, 2017). Most theories of value-based decision-making have focused on processes related to the incremental learning of value following external reinforcement, but have not explicitly addressed the role of memory per se. Thus, fundamental questions remain regarding interactions between memory and value-based decisions, which have been gaining attention in recent years.

Several recent empirical studies have demonstrated interactions between episodic memory and value-based decision-making. Memory for past events has been shown to bias value-based decisions (K. D. Duncan & Shohamy, 2016), differently for choices of novel versus choices of familiar options (K. Duncan, Semmler, & Shohamy, 2019). Memory accessibility has been shown to affect people’s risk preferences (Kusev, van Schaik, & Aldrovandi, 2012). Choice behavior and fMRI signals during value-based decision-making were better explained by episodic memory for individual past choices than by a standard reinforcement learning model (Bornstein, Khaw, Shohamy, & Daw, 2017). Another study has found that during sampling of episodic memories of previous choices, the retrieved context influenced present choices, deviating from the predictions of standard reinforcement learning models (Bornstein & Norman, 2017). Furthermore, it has been shown that choices are influenced by value that is spread across memories from rewarded stimuli to associated items that were never experienced directly with reward (Wimmer & Shohamy, 2012). At the neural level, the ventromedial prefrontal cortex (vmPFC) and the hippocampus both have been shown to play a role in memory processes and value-based decisions (Weilbächer & Gluth, 2017) and recent studies have been further emphasizing that the hippocampus bridges between past experience and future decisions (Bakkour et al., 2019; Biderman, Bakkour, & Shohamy, 2020).

All these studies highlighted the interaction between memory and value-based decision-making involving external reinforcements. However, everyday life involves decisions and associations that are not directly reinforced. Thus, it remains unclear if memory plays a general role in value-based decision-making even without external reinforcements.

To better understand the role of memory processes in shaping preferences independently of external reinforcements, we utilized a novel behavioral change paradigm, named cue-approach training (CAT). In this paradigm, associating images of items with a neutral cue and a speeded motor response results in a consistent preference enhancement without external reinforcement which is maintained for months (Bakkour et al., 2018; Botvinik-Nezer, Salomon, & Schonberg, 2019; Salomon et al., 2018; Salomon, Botvinik-Nezer, Oren, & Schonberg, 2019; Schonberg et al., 2014). During CAT, images of items are consistently paired with a neutral cue and a speeded motor response (*‘Go items’*), while other items are presented without the cue nor the response (*‘NoGo items’*). One training session with several presentations of all items leads to long-lasting preference changes, measured as the likelihood of choosing Go over NoGo items that had similar initial subjective values (Schonberg et al., 2014). Results from over 30 samples with this paradigm have demonstrated a replicable effect on various types of stimuli, including snack food items, fruits and vegetables, unfamiliar faces, fractal art images and positive affective images (Bakkour et al., 2018, 2016; Bakkour, Lewis-Peacock, Poldrack, & Schonberg, 2017; Botvinik-Nezer et al., 2019; Salomon et al., 2018, 2019; Veling et al., 2017; Zoltak, Veling, Chen, & Holland, 2017), revealing the potential of the CAT paradigm as an experimental platform for value-based decision-making without external reinforcements (Schonberg & Katz, 2020).

The underlying mechanisms of the change of preferences following CAT are not yet fully understood (Bakkour et al., 2017; Botvinik-Nezer et al., 2019; Salomon et al., 2019; Schonberg et al., 2014; Schonberg & Katz, 2020). The long-lasting nature of the effect, which has been shown to last for up to six months following a single training session (Botvinik-Nezer et al., 2019; Salomon et al., 2018, 2019; Schonberg et al., 2014), raises the hypothesis that memory processes are involved in its maintenance. Preferences as memories (PAM) is a framework that suggests that the retrieval of relevant knowledge from memory is the basis used to compare options and make a choice between them (Weber & Johnson, 2006). This framework provides an explanation for the associative nature of preferences, as well as known characteristics of preferences such as instability and semantic framing-effects. Based on this framework, we hypothesized that CAT enhances memory of Go items, which in turn leads to preferring these items over NoGo items. Previous neuroimaging findings with CAT that suggested possible interactions between hippocampal fMRI activity and subsequent preferences one month following CAT, provide additional evidence in support of this hypothesis (Botvinik-Nezer et al., 2019). Therefore, here we set out to directly test, for the first time, the role memory processes play in the behavioral change of preferences following CAT, in the short and in the long-term.

We propose an underlying mechanism for the CAT effect, in which preference change following CAT results from a boost in memory encoding of positive Go items, which in itself is a consequence of enhanced perceptual processing of Go items (Botvinik-Nezer et al., 2019; Schonberg et al., 2014). We hypothesize that the enhanced encoding of Go items, as well as the greater perceptual activation in response to them, increases accessibility of attributes and associations of these specific Go items (Anderson, 1983; Bhatia, 2013).

In order to test memory for individual items, in the current work we introduced a memory recognition task following CAT. In two independent pre-registered experiments and one pilot experiment, memory was evaluated following a long (16 repetitions) or short (a single exposure) CAT training session, before the probe phase that evaluated post-training preferences. We then tested (1) whether memory is stronger for Go compared to NoGo items following CAT and (2) whether memory is related to choices.

We pre-registered our predictions that responses in the memory recognition task will be both faster and more accurate for Go compared to NoGo items. Furthermore, we hypothesized that preference changes, reflected in the binary choice phase, are due to the enhanced accessibility of memory associations of the Go items, which tips the scales in favor of the Go items. Therefore, we pre-registered our prediction that better remembered Go items will be chosen over worse remembered NoGo items. These hypotheses were tested both in the short-term (immediately or a few days after CAT) and in a one-month follow-up.

## Experiment 1

In the first pre-registered experiment, we tested whether memory was enhanced for Go compared to NoGo items following CAT and whether memory was related to choices. This experiment included three sessions: Session 2 was completed exactly three days after Session 1 and tested memory involvement in the (relatively) short-term effect of CAT. The Follow-up Session, testing the long-term effects, was performed about one month following Session 1.

Thirty-nine participants completed the first two sessions. Four participants were excluded based on pre-registered exclusion criteria. Thirty participants also completed the Follow-up Session, about one month after Session 1 (mean = 34.3 days, *SD* = 5.04 days, range = 28 - 46 days; demographic statistics are reported in Table 1).

**Table 1.**
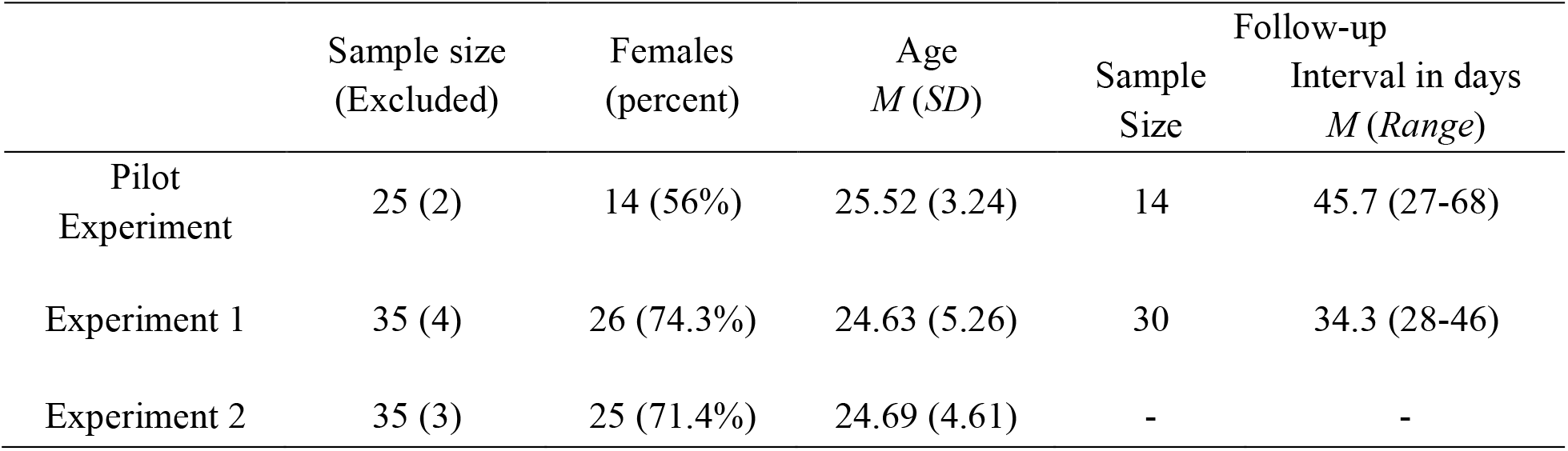
*Demographic Information*

In Session 1, we first evaluated participants’ preferences for 60 snack food items with a Becker-DeGroot-Marschak (BDM) auction procedure (Becker, DeGroot, & Marschak, 1964) (Figure 1a). Then, they completed 16 repetitions of CAT with 40 of the 60 snack food items (Figure 1b). In Session 2, we tested participants’ memory with a recognition task (Figure 1c) that included 28 new and 28 old (i.e., previously presented) snack food items. We then tested whether their preferences were modified with a binary forced choice probe task (Figure 1d; see Supplementary Table 1 for item allocation to value categories), and finally an additional BDM auction (Figure 1a). The Follow-up Session included the same tasks as in Session 2.

**Figure 1.**
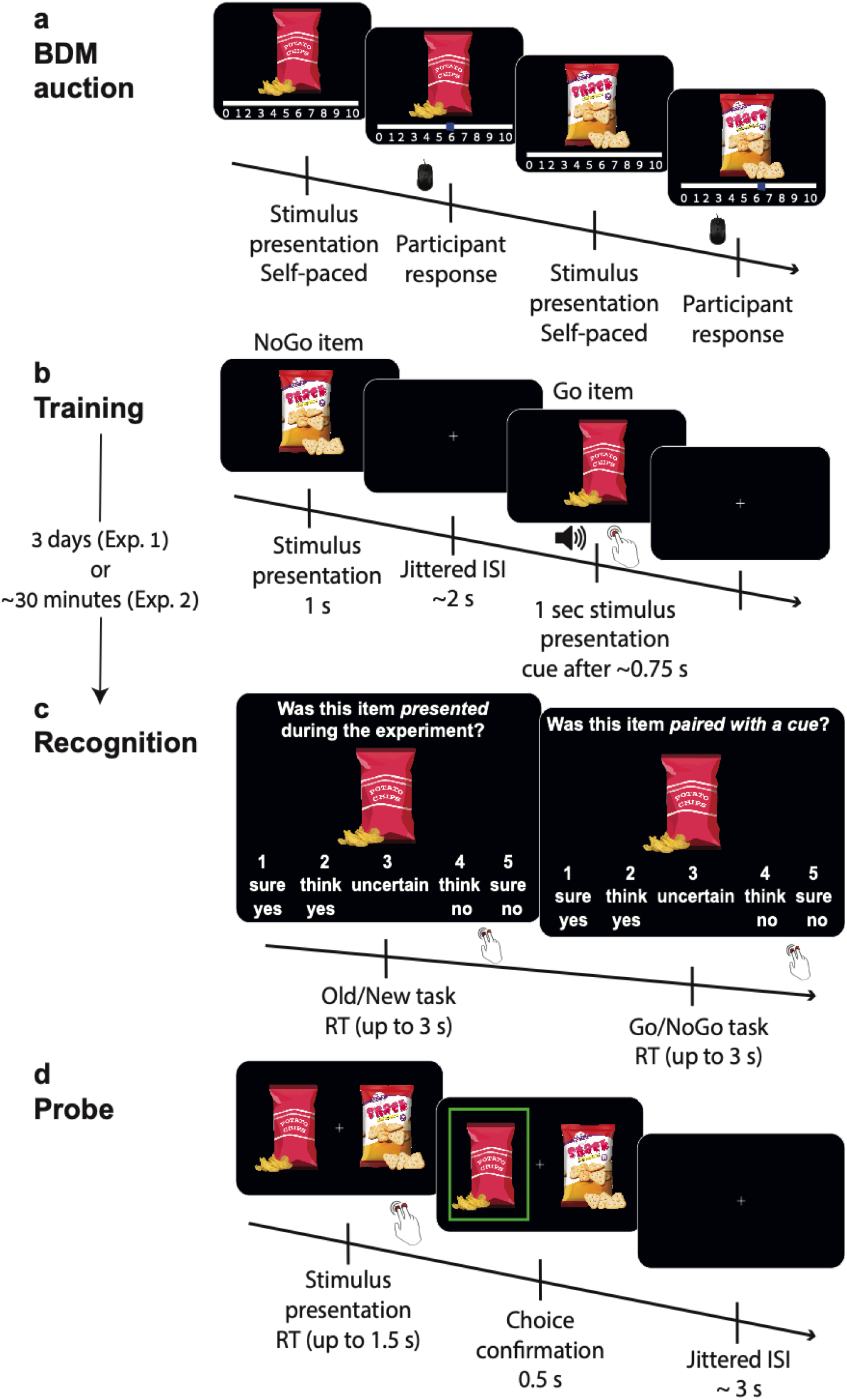
Outline of the experimental procedures. (a) Initial preferences were evaluated using the Becker-Degroot-Marschak (BDM) auction procedure. (b) Cue-approach training (CAT): Go items were associated with a neutral auditory cue and a speeded response (16 repetitions per stimulus in Experiment 1, and only one repetition per stimulus in experiment 2). (c) Recognition memory task. (d) Probe: Participants chose between pairs of Go and NoGo items of similar initial value. In Experiment 1, the recognition and probe tasks were completed in Session 2, three days following the first session, and then repeated again in the Follow-up Session, about one month following the first session. In Experiment 2, all tasks were performed on the same day, and a 30-minutes filler task separated between the training and the recognition task.

## Results

### Recognition

Overall, participants largely showed the pre-registered predicted pattern of behavior in both sessions: participants better remembered Go compared to NoGo items and there was no interaction with the value category.

#### *Session 2.* (for full statistics see Supplementary Table 2)

##### Overall performance

The mean hit rate of participants was 86.31% (*SD* = 10.33%). The mean correct rejection rate was 70.53% (*SD* = 17.60%). The mean d’ was 1.824 (*SD* = 0.747). The mean RT was 1.458 (*SD* = 0.262) seconds for hits, 1.796 (*SD* = 0.504) seconds for misses, 1.589 (*SD* = 0.351) seconds for correct rejections and 1.840 (*SD* = 0.366) seconds for false alarms.

##### Hit rate

Overall, as predicted and pre-registered, hit rate in the old / new recognition task was significantly higher for Go (*M =* 88.51%) compared to NoGo (*M =* 84.97%) items (*one-sided p* = .008, *odds ratio* = 2.873, 95% *CI* [1.225, 6.737], mixed-effects logistic regression; see Figure 2a), while it was not higher for high-value (*M* = 86.30%) compared to low-value (*M* = 87.40%) items (*p* = .437). No interaction was found between the value category (high-value / low-value) and item type (Go / NoGo; *two-sided p* = .472).

**Figure 2.**
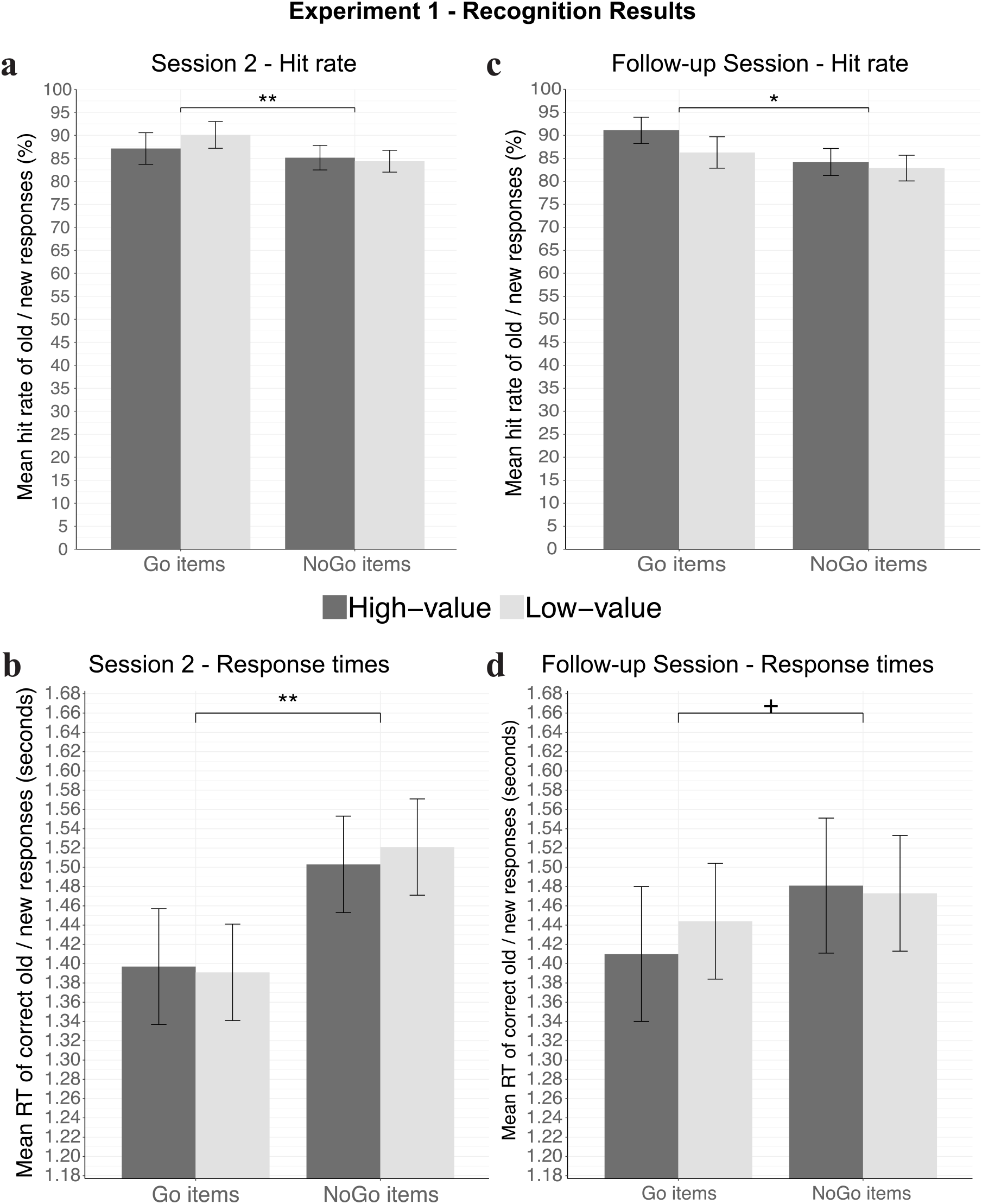
Recognition results of Experiment 1. The mean hit rate (a,c) and response times (b,d) of each participant were calculated and then averaged across participants. Error bars represent standard error of the mean across participants. Asterix represent one-sided statistical significance; + *p* < .07, * *p* < .05, ** *p* < .01

##### Response times

In accordance with our pre-registered predictions, RTs in the recognition task were significantly faster for Go (*M* = 1.392 seconds) compared to NoGo (*M* = 1.514 seconds) items (*one-sided p* = .001, *estimated mean difference* = -0.124, mixed-effects linear regression; see Figure 2b). We did not find RT differences between high-value (*M* = 1.448 seconds) and low-value (*M* = 1.454 seconds) items (*one-sided p* = .435, *estimated mean difference* = -0.005) nor an interaction between the value category and item type (*two-sided p* = .577).

##### Follow-up Session

(for full statistics see Supplementary Table 2). In this session, we tested the relationships between memory and choices in the long-term, one month following CAT.

##### Overall performance

The mean hit rate of participants was 84.47% (*SD* = 11.76%). The mean correct rejection rate was 61.15% (*SD* = 19.85%). The mean d’ was 1.454 (*SD* = 0.742). The mean RT was 1.456 (*SD* = 0.314) seconds for hits, 1.633 (*SD* = 0.446) seconds for misses, 1.603 (*SD* = 0.326) seconds for correct rejections and 1.749 (*SD* = 0.336) seconds for false alarms.

##### Hit rate

As in Session 2 and in accordance with our pre-registered predictions, hit rate in the old / new recognition task was significantly higher for Go (*M* = 88.81%) compared to NoGo (*M* = 83.53%) items (*one-sided p* = .017, *odds ratio* = 1.981, 95% *CI* [1.055, 3.718], mixed-effects logistic regression; see Figure 2c) and not significantly higher for high-value (*M* = 87.57%) compared to low-value (*M* = 84.77%) items (*one-sided p* = .123, *odds ratio* = 1.379, 95% *CI* [0.788, 2.412]). As in Session 2, there was no interaction between the value category and item type (*two-sided p* = .394).

##### Response times

A general trend of faster RTs for Go (*M* = 1.422 seconds) compared to NoGo (*M* = 1.477 seconds) items was observed, but only with marginal significance (*one-sided p* = .063, *estimated mean difference* = -0.049, mixed-effects linear regression; see Figure 2d). RTs were not significantly faster for high-value (*M* = 1.442 seconds) compared to low-value (*M* = 1.455) items (*one-sided p* = .314, *estimated mean difference* = -0.015). There was no interaction between the value category and item type (*p* = .636).

#### Choices - all sessions

(see Figure 3; for full statistics see Supplementary Table 3). As predicted and pre-registered, replicating previous findings with CAT (Botvinik-Nezer et al., 2019; Salomon et al., 2018; Schonberg et al., 2014), participants significantly chose high-value Go over high-value NoGo items, both three days after CAT (*M* = 61.67%, *one-sided p* < .001, *odds ratio* = 1.745, 95% *CI* [1.647, 1.849], mixed-effects logistic regression) and one month after CAT (*M* = 59.62%, *one-sided p* < .001, *odds ratio* = 1.546, 95% *CI* [1.464, 1.632]). They also significantly chose low-value Go over low-value NoGo items three days after CAT (*M* = 58.16%, *one-sided p* = .009, *odds ratio* = 1.487, 95% *CI* [1.387, 1.594]) and marginally in the one-month follow-up (*M* = 54.95%, *one-sided p* = .066, *odds ratio* = 1.244, 95% *CI* [1.164, 1.330]). Unlike previous experiments with snack food items, the proportion of Go item choices was not significantly higher for high-value compared to low-value probe pairs in either one of the two sessions. However, it should be noted that we improved the models used in previous studies, which is likely the reason for the lack of replication.

**Figure 3.**
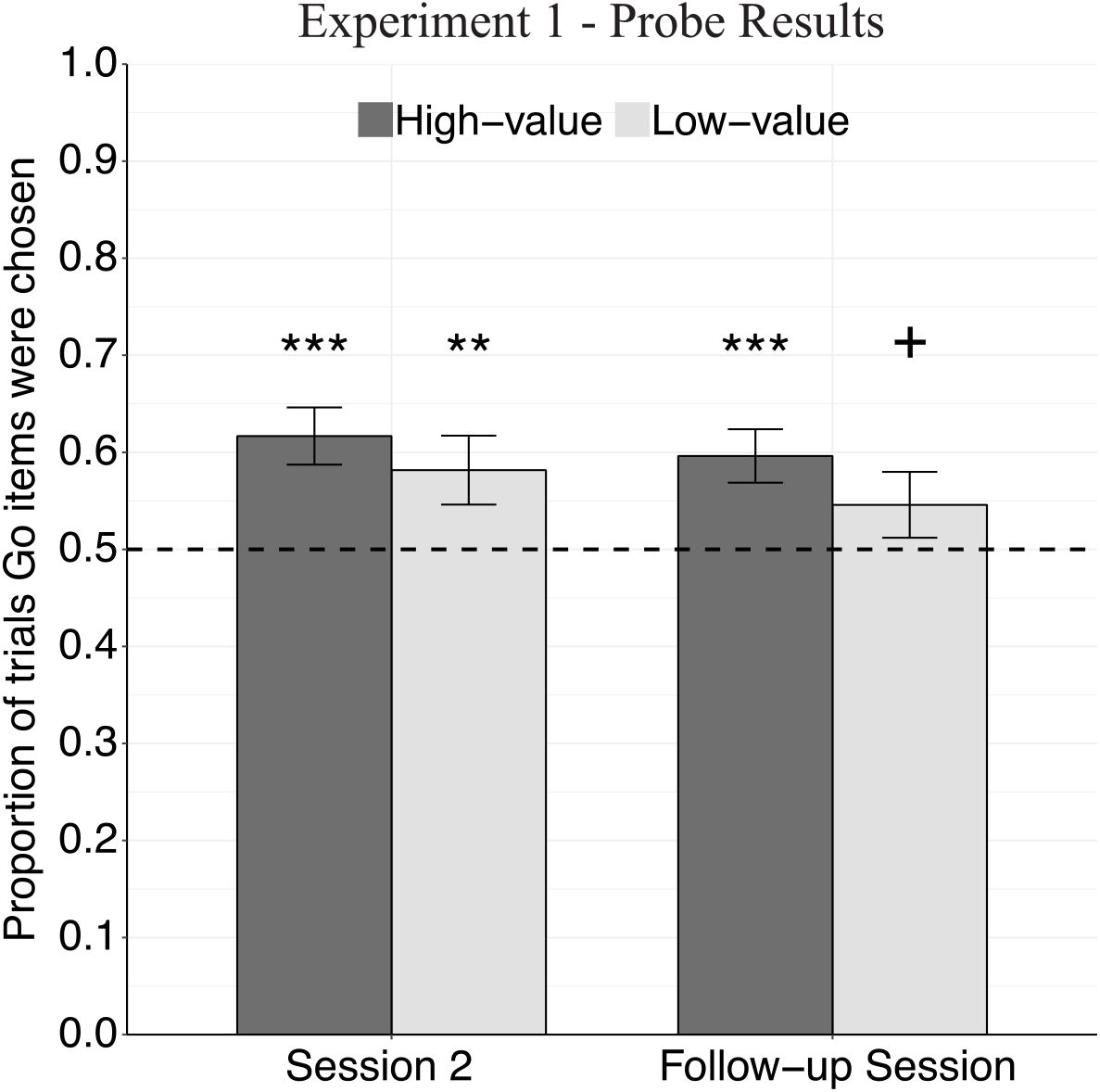
Probe results of Experiment 1. Mean proportions of choosing the Go item (calculated for each participant and then averaged across participants) are presented with error bars representing standard error of the mean. Dashed line indicates 50% chance level. Asterisks represent statistical significance of a one-sided logistic regression analysis; + *p* < .07, * *p* < .05, ** *p* < .01, *** *p* < .001.

#### Relationships between memory and choices

Our main analysis focused on the relationships between memory and choices at the individual item level.

#### *Session 2.*(see Supplementary Table 4)

We tested the relationship between memory for the specific items and choices made in the subsequent probe in Session 2. Based on our suggested underlying mechanism of CAT, we predicted that these relationships will be significantly positive for choices between high-value items. We further predicted a significant interaction between memory and the pair value category (high-value / low-value).

Participants chose Go items significantly more when the Go item was remembered and the NoGo item was forgotten, compared to when both were remembered or forgotten (see Figure 4a), both for choices between high-value items (*one-sided p* = .026, *odds ratio* = 2.258, 95% *CI* [0.993, 5.135], mixed-effects logistic regression) and for choices between low-value items (*one-sided p* = .024, *odds ratio* = 1.341, 95% *CI* [1.003, 1.792]), with a significant interaction between this accuracy category and the value category (*one-sided p = .040)*. There were no significant differences in Go item choices when comparing pairs in which the Go item was forgotten and the NoGo item was remembered, to pairs in which both items were remembered or forgotten, neither for choices between high-value items (*one-sided p* = .102) nor for choices between low-value items (*one-sided p* = .893). The interaction effect was also not significant (*one-sided p* = .088).

**Figure 4.**
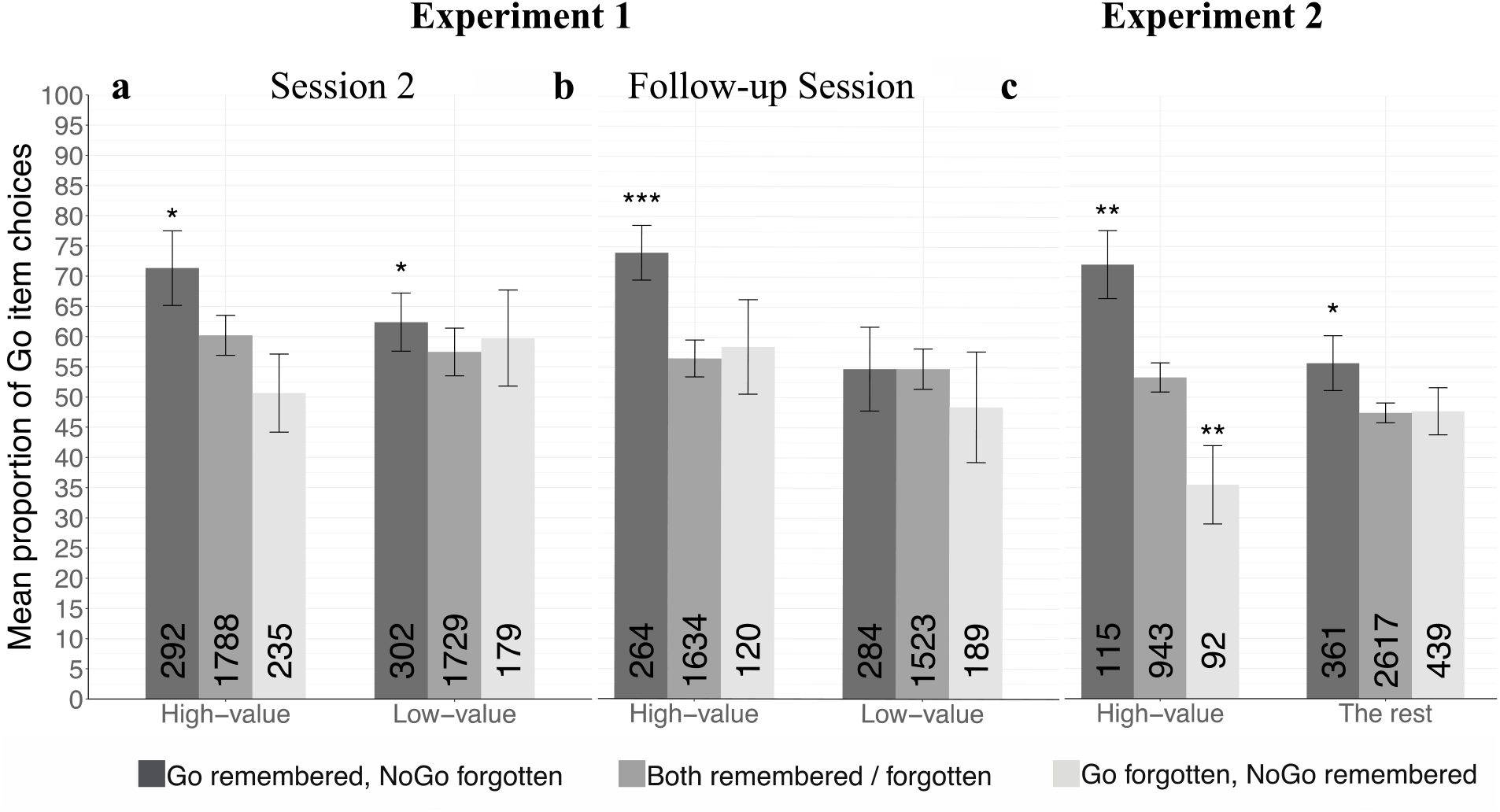
The relationships between recognition memory and choices, in (a) Experiment 1, Session 2 (performed three days after Session 1); (b) Experiment 1, Follow-up Session (∼30 days after Session 1); (c) Experiment 2. Mean percent of Go item choices as a function of recognition memory with error bars representing standard error of the mean. The number of trials, summed across participants, is presented at the bottom of each bar. Asterisks represent statistical significance between each category and the “Both remembered / forgotten” baseline category (one-sided mixed-effects logistic regression); * *p* < .05, ** *p* < .01, *** *p* < .001.

The relationship between the recognition ΔRT and choices (see Figure 5a) of Go over NoGo items was not significant, neither for high-value items (*one-sided p* = .171, mixed-effects logistic regression) nor for low-value items (*one-sided p* = .302).

**Figure 5.**
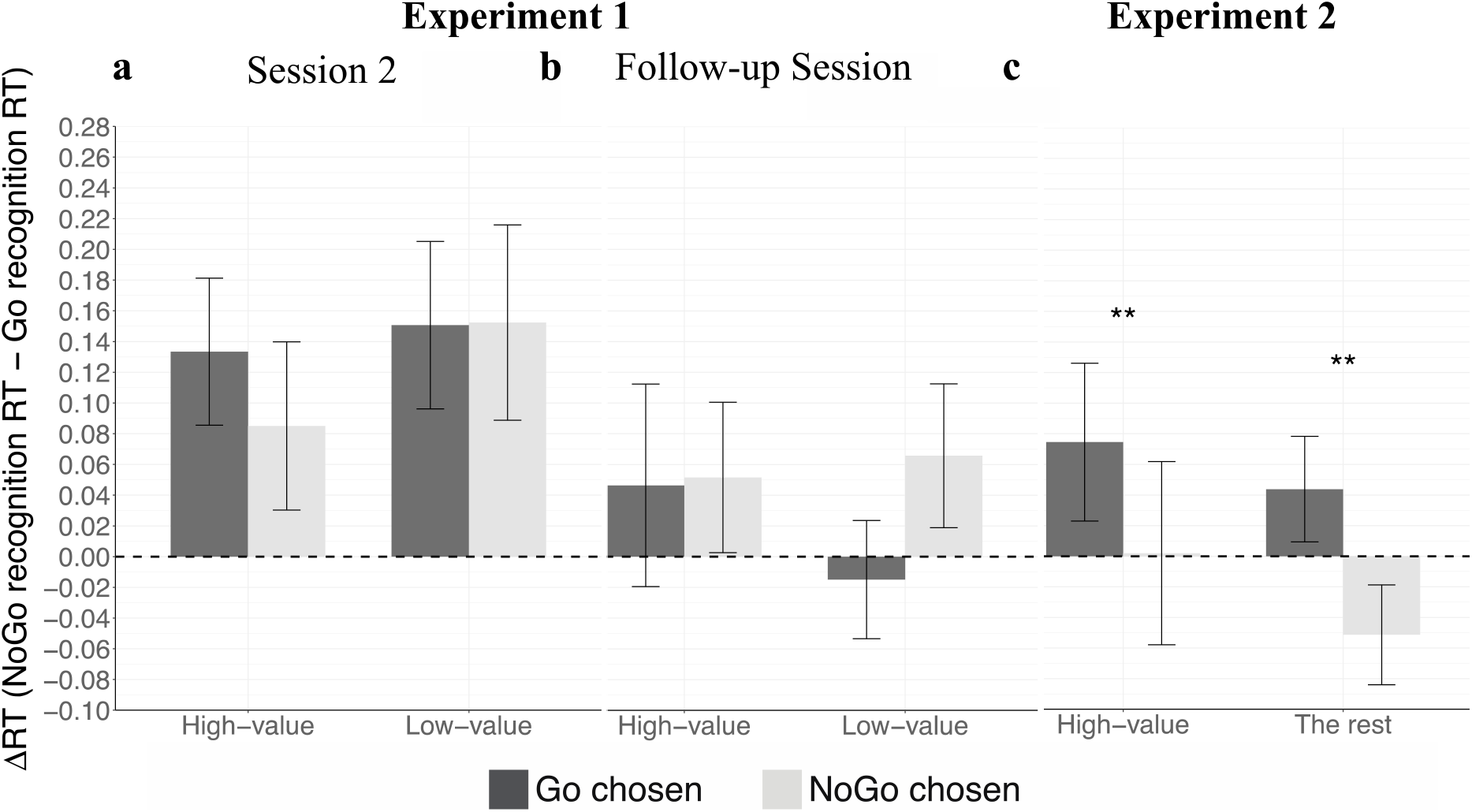
The relationships between recognition ΔRT and choices in (a) Experiment 1, Session 2 (performed three days after Session 1); (b) Experiment 1, Follow-up Session (performed ∼30 days after Session 1); (c) Experiment 2. Mean ΔRT values (in seconds) are presented for choices of Go items and for choices of NoGo items, within each value category. Error bars represent standard error of the mean. Asterisks represent statistical significance of one-sided mixed-effects logistic regression; ** *p* < .01

#### Follow-up Session

(see Supplementary Table 5). Participants chose Go items more when the Go item was remembered and the NoGo item was forgotten, compared to when both were remembered or forgotten (see Figure 4b), when choosing between high-value items (*one-sided p* < .001, *odds ratio* = 2.810, 95% *CI* [1.774, 4.450], mixed-effects logistic regression), but not when choosing between low-value items (*one-sided p* = .237), with a significant interaction between the accuracy category and the value category (*one-sided p* = .002, meaning that this effect was significantly stronger for high-value compared to low-value items). Again, there were no significant differences in Go item choices when comparing pairs in which the Go item was forgotten and the NoGo item was remembered, to pairs in which both items were remembered or forgotten, neither for choices between high-value items (*one-sided p* = .589, mixed-effects logistic regression) nor for choices between low-value items (*one-sided p* = .503). The interaction effect was also not significant (*one-sided p* = .454). The relationship between the ΔRT and choices (see Figure 5b) of Go over NoGo items was not significant, neither for high-value items (*one-sided p* = .174, mixed-effects logistic regression) nor for low-value items (here the direction of the difference was negative: when the ΔRT between the low-value NoGo and the low-value Go item was larger, the Go item was less likely to be chosen; *one-sided p* = .924). The interaction between the ΔRT and value category was significant (*one-sided p* = .005).

## Discussion

In Experiment 1, similar to previous studies, we found enhanced preferences towards Go items, both three days and one month following CAT. Unlike previous studies with snack food items (Botvinik-Nezer et al., 2019; Schonberg et al., 2014), the CAT effect was not significantly stronger in high-value than in low-value probe choices in this experiment (Figure 3). This is likely a result of using different statistical model (i.e., modeling both the random intercept and random slope).

In line with our pre-registered suggested mechanism, we found enhanced memory for Go compared to NoGo items (reflected as higher hit rate and faster RTs), both in the short- and in the long-term. This is the first direct evidence for enhanced memory accessibility of Go versus NoGo items following CAT, suggesting that memory modifications are involved in the underlying mechanism of the CAT effect.

Most importantly, since we hypothesized that each choice is related to the memory for the specific items in the presented pair, we tested whether choices were related to the memory for the specific items on each binary choice pair. We found evidence showing that memory was indeed related to the specific choices. Choices of the Go items over the NoGo items were more likely when the individual Go item was remembered and the NoGo item was forgotten. Furthermore, these relationships were generally more positive for choices between high-value items compared to choices between low-value items. These results are in line with our hypothesis, that for higher valued items, better remembered Go items will be chosen over worse remembered NoGo items.

It should be noted that we originally planned to test recognition memory with paired t-tests and the relationships between memory and choices with linear correlation tests (see deviation from pre-registration section). The results of the t-tests are similar to the results of our reported results and are mostly significant (see Supplementary Materials: Pre-Registered Analyses for Experiment 1). The results of the linear correlation tests are mostly insignificant. However, these linear correlations test the relationships between memory modifications and choices across participants, rather than across items, and are less suited compared to the tests we report here. The fact that correlation tests across participants yielded mostly insignificant results, while correlations across items yielded mostly significant results, further emphasize that CAT has an item-specific rather than an overall effect on items.

Experiment 1 was the first experiment with CAT in which choices were first tested three days (and not immediately) after training. In previous experiments there was always a probe phase on the same day of training and then subsequent probes up to six months after the initial training. The results of Experiment 1 show that training directly influences choices for at least a few days after training, therefore suggesting that the long-term duration of the CAT effect is less likely to be driven by previous choices via a mere choice effect (Sharot, Fleming, Yu, Koster, & Dolan, 2012).

## Experiment 2

The findings of Experiment 1 suggested that choices following CAT were related to memory at the individual item level. To further examine these relationships and replicate the effect we found, in Experiment 2, we sought to increase the variance of memory scores across items and allow a better examination of the memory effects on individual items. To do so, we doubled the number of trained and tested items, and for the first time, we decreased the number of training repetitions to one. Our goal was to increase the variance across items, in pre-training preferences, post-training memory and post-training choices, to further examine the relationships between memory and value-based choices at the individual item level.

N = 35 valid participants completed Experiment 2, which was also pre-registered. Three additional participants were excluded based on our pre-registered exclusion criteria. Upon arrival, participants completed a BDM auction with 80 items (instead of 60 items as in Experiment 1), Then, they completed the training task, which was similar to Experiment 1, besides two important modifications: First, each item was only presented once (instead of 16 repetitions in Experiment 1). Second, 80 (instead of 40) items were presented during training, with 12 high-value Go items and 12 low-value Go items (compared to six of each category in Experiment 1). Following the training task, participants completed a filler task that lasted about 30 minutes. Then, their memory was tested with a recognition task, similar to the one performed in Experiment 1, but with a much larger total number of 160 items - 80 old and 80 new items. Finally, participants’ preferences were revaluated with the probe task that included a doubled number of choices compared to Experiment 1.

## Results

### Recognition

(see Supplementary Table 6). In contrast to our predictions, participants did not remember the Go compared to the NoGo items more accurately following one training run.

#### Overall performance

The mean hit rate of participants was 85.63% (*SD* = 11.34%). The mean correct rejection rate was 81.6% (*SD* = 12.48%). The mean d’ was 2.403 (*SD* = 1.829). The mean RT was 1.384 (*SD* = 0.214) seconds for hits, 1.558 (*SD* = 0.290) seconds for misses, 1.436 (*SD* = 0.239) seconds for correct rejections and 1.752 (*SD* = 0.349) seconds for false alarms.

#### Hit Rate

In contrast to Experiment 1 and to our pre-registered prediction, in Experiment 2 the hit rate in the old / new recognition task was not higher for Go (*M* = 84.67%) compared to NoGo (*M* = 85.78%) items (*one-sided p* = .759, mixed-effects logistic regression; see Supplementary Figure 1a), but was significantly higher for higher value categories (*one-sided p* = .004). There was no interaction between the value category (high / medium-high / medium-low / low value) and item type (Go / NoGo; *two-sided p* = .570).

#### Response times

Contrary to our predictions, RTs were not significantly faster for Go (*M* = 1.380 seconds) compared to NoGo (*M* = 1.383 seconds) items (*one-sided p* = .484, mixed-effects linear regression; see Supplementary Figure 1b), nor significantly faster for higher value items (*one-sided p* = .211). The interaction between item type and value category was not significant (*two-sided p* = .738).

### Choices

(for full statistics see Supplementary Table 7). In the high-value probe choices (equivalent to those of Experiment 1), participants significantly chose Go over NoGo items (*M* = 54.41%, *one-sided p* = .035, *odds ratio* = 1.209, 95% *CI* [1.152, 1.270], mixed-effects logistic regression; see Supplementary Figure 2) after one training run. However, participants did not show preference modification for the Go items in the other value categories, nor when combining the high and medium-high value categories to one category.

### Relationships between memory and choices (see Supplementary Table 8)

In Experiment 2, we tested the relationships between choices and memory within each of the four value categories. Since preferences were only changed for high-value items, we combined the other three categories (i.e., “The rest”. See Supplementary Table 9 for statistics of each of these value categories) in our analysis. We tested the interactions between memory and value category by modelling the value category as high-value compared to the rest of the categories. Participants chose Go items significantly more when the Go item was remembered and the NoGo item was forgotten compared to when both were remembered or forgotten (see Figure 4c), both for choices between high-value items (*one-sided p* = .004, *odds ratio* = 2.627, 95% *CI* [1.275, 5.410], mixed-effects logistic regression) and for choices between items in the rest of the value categories combined (*one-sided p* = .016, *odds ratio* = 1.287, 95% *CI* [1.023, 1.619]). The interaction between this accuracy category and the value category (high-value compared to the rest) was significant (*one-sided p* = .016). Participants chose Go items significantly less when the Go item was forgotten and the NoGo item was remembered, compared to when both were remembered or forgotten (see Figure 4c), when choosing between high-value items (*one-sided p* = .009, *odds ratio* = 0.499, 95% *CI* [0.280, 0.888], mixed-effects logistic regression), but not when choosing between items from the rest of the value categories (*one-sided p* = .270). The effect was significantly stronger for choices between high-value items compared to choices between items from the rest of the value categories (an interaction effect; *one-sided p* = .009).

The relationship between the ΔRT and choices of Go over NoGo items (see Figure 5c) was significantly positive for the high-value items (*one-sided p* = .004, *odds ratio* = 1.511, 95% *CI* [1.108, 2.059], mixed-effects logistic regression) as well as for the rest of the value categories combined (*one-sided p* = .002, *odds ratio* = 1.371, 95% *CI* [1.101, 1.708]). The interaction between the ΔRT and value category was not significant (*one-sided p* = .329).

## Discussion

In Experiment 2, we tested the relationship between choices and memory for individual items with a larger number of items compared to previous experiments and following one repetition of CAT. Even after a single training run, we found a preference modification effect for items in the high-value category. We did not, however, find such effect in the other value categories. Contrary to our pre-registered predictions, we found no evidence for a main effect of better memory of Go items in comparison to NoGo items. These results imply that one training run might be sufficient to change preferences for the most valued Go items (a change which was significant but weak compared to previous CAT samples), but not sufficient to induce an effect of enhanced memory across items.

Replicating the results of Experiment 1, and in line with our pre-registered hypotheses, we found strong relationships between better memory for the Go items and choices of these items over the less remembered NoGo items, mainly for high-value items (the value category in which the choice effect was found). The larger number of Go and NoGo items used in the recognition and probe tasks in this experiment compared to previous ones (48 items in the current experiment versus 24 or less in previous experiments), allowed us to more generally test the relationships between memory for specific items and choices. These results provide additional evidence in support of our suggested mechanism, by replicating, in a different design, the finding that better-remembered Go items following CAT are also chosen more often.

## General Discussion

The role of memory processes in value-based decision-making, as well as the cognitive and neural mechanisms by which memory influences decisions, are not yet fully understood (Biderman et al., 2020; Fellows, 2017; Schonberg & Katz, 2020; Shadlen & Shohamy, 2016; Weilbächer & Gluth, 2017). Recent studies have begun to highlight the interactions between memory and value-based decision-making. However, most of these studies involved external reinforcements, while everyday life involves decisions and associations that are not directly reinforced (Schonberg & Katz, 2020).

Here, we tested the role memory processes play in the behavioral change of preferences following the CAT paradigm. This paradigm has been shown to reliably induce long-lasting preference change using the mere association of images of items with a neutral cue and a speeded button press response, without external reinforcements (Botvinik-Nezer et al., 2019; Salomon et al., 2018, 2019; Schonberg et al., 2014). We hypothesized that the cognitive mechanisms underlying the CAT effect are based on modifications of memory accessibility. We suggested that memory for Go items is enhanced following CAT. Therefore, we predicted better recognition memory for Go compared to NoGo items following CAT in the short and in the long-term, and that the better memory of Go items lead to greater preferences for high-value Go items during probe.

We tested our predictions in two pre-registered experiments based on an additional pilot experiment. In Experiment 1, we tested these predictions with a recognition and a subsequent probe task, performed three days after CAT. Similar to previous studies with CAT (Bakkour et al., 2018, 2016, 2017; Botvinik-Nezer et al., 2019; Salomon et al., 2018, 2019; Schonberg et al., 2014; Veling et al., 2017; Zoltak et al., 2017), we found enhanced preference for Go items following CAT. High-value Go items were preferred over high-value NoGo items three days as well as one month following CAT (Figure 3). Low-value Go items were also preferred over low-value NoGo items. As our suggested mechanism predicted, we also found a significant memory enhancement, reflected as higher hit rates and faster RTs in the recognition task for Go compared to NoGo items, both in the short and in the long-term (Figure 2). Moreover, we found that memory for individual items was positively related to choices of Go over NoGo items, both when memory and choices were tested three days as well as one-month after CAT, mainly for choices between high-value items (Figures 4 and 5).

The findings of Experiment 1 suggested that CAT affects memory, and thus choices, at the individual item level. Therefore, in Experiment 2, we increased the number of items to more widely explore the variability of the effect across items. We decreased the number of training repetitions to one, in order to test whether one run is sufficient to affect memory and preferences. We found that even a single training run was sufficient to induce preference modification in the highest value category tested. Overall, memory was not significantly enhanced for Go compared to NoGo items. Nonetheless, we found significant relationships between memory and choices: better remembered Go items were more likely to be preferred over worse remembered NoGo items in the binary probe phase, mainly for choices between high-value items. The absence of overall memory modifications might have been the result of lack of consolidation in such short time scales (memory was tested about 30 minutes after CAT).

Prior to these experiments we conducted a pilot experiment (see Supplementary Materials), which was similar to Experiment 1, but with an additional probe task performed at the end of Session 1. In this pilot sample we also found positive relationships between memory for individual Go items and choices of these items over worse remembered NoGo items. Unlike previous studies, there was no overall CAT effect in the pilot experiment. We also did not find enhanced memory for Go compared to NoGo items, possibly due to a ceiling effect in recognition memory performance in this experiment. Nevertheless, the existence of a link between memory and choices at the individual item level in the Pilot Experiment indicates that even when the main effects of both choices and memory are absent, individual choices are still related to memory for the individual items.

We interpreted faster RTs for Go compared to NoGo items in Experiment 1 as an indication of better memory for Go items, in accordance with our pre-registered hypotheses and predictions. However, an alternative explanation for this finding might be that the association of Go items with motor responses during training resulted in a conditioned response of pressing when Go items are presented, leading to faster RTs for these items irrespective of memory strength. To disentangle these two hypotheses, we performed exploratory analyses on the recognition data from Experiment 1 (see Supplementary Exploratory Analyses). The results of this analysis suggest that the faster RTs for Go compared to NoGo items in the old / new recognition task are indeed an indication of better memory for these items, and not of a response bias.

Response bias can also putatively explain the interactions we found between recognition memory and choices following CAT, leading participants to respond affirmatively to Go items in both the memory recognition and the binary-choice probe tasks. However, findings of the current as well as previous studies do not support this possibility. First, CAT affects choices between high-value items but frequently not choices between low-value items (Botvinik-Nezer et al., 2019; Schonberg et al., 2014). A response bias would have been predicted to also affect low-value choices. Moreover, the CAT effect was also found in a previous sample when choices were made with the eyes rather than the hands (Bakkour et al., 2016), further indicating that the effect of CAT is not solely a response bias in the hand motor circle.

Another alternative explanation for the interactions we found between memory and choices following CAT, is that CAT influences a third variable, such as familiarity, which affects both choices and memory. However, all items included in our stimuli dataset are local familiar snack food items, and our item allocation procedure, which matches choice pairs based on the initial WTP, ensures that any pre-CAT difference between Go and NoGo items would be random. Furthermore, the number of times each item is presented during CAT is the same for Go and NoGo items, therefore a mere exposure effect cannot explain the CAT effect (Zajonc, 1968). Nevertheless, the current study did not explicitly manipulate familiarity, and future studies can test this alternative explanation, for example by comparing familiar to unfamiliar items (e.g., local snacks versus snacks from other countries, or familiar versus unfamiliar faces). Another third variable might be emotional devaluation towards NoGo items as a result of inhibition (Driscoll, Launay, & Fenske, 2018; Fenske & Raymond, 2006). However, it is not clear whether emotional devaluation would lead to the observed memory differences and correlations with choices. This can also be directly tested in future studies.

An interesting aspect of CAT with snack food items is that it affects choices of high-value Go over NoGo items more than choices of low-value Go over NoGo items (Botvinik-Nezer et al., 2019; Salomon et al., 2018; Schonberg et al., 2014). This effect was suggested to indicate that CAT may induce value scaling, rather than a fixed-increment value enhancement (McGuire & Kable, 2014). Alternatively, we suggest that this differential effect is due to the greater number of positive memories associated with high-value versus low-value snacks. Our proposed mechanism suggests that memory is enhanced for both high-value and low-value Go items following CAT. When presented with binary choices of high-value items, the memories associated with Go items that are activated are usually positive, resulting in a preference for these items. When choosing between low-value items, activated associations are not necessarily positive since these items are subjectively less preferred to begin with (but are still mostly liked). Therefore, choices of low-value Go over low-value NoGo items following CAT are significantly less consistent.

Unlike previous studies with CAT and snack food items, here we did not find a significant differential effect, in which the proportion of Go items choices is higher for high-value versus low-value items. However, it should be noted that in the current study we used a different and putatively improved random effects structure. Results were more similar to previous studies when using the same models we used in our older work. This suggests that the variance was likely underestimated in some of the previous results and explains why we did not replicate some of the previous findings in this work, with the new mixed-effects models.

Overall, the findings of the current study were in line with some of our pre-registered hypotheses. When choices were affected, memory was enhanced for Go compared to NoGo items. Memory was enhanced for Go compared to NoGo items, irrespective of value category (low-value or high-value), suggesting that there is no differential effect of CAT on memory (i.e., that memory enhancement is not stronger for high-value Go compared to low-value Go items). Most importantly, when examining memory for each of the items in individual binary choices, we found that better memory for the Go compared to the NoGo items was related to choices of Go over NoGo items more positively for high-value compared to low-value choice pairs.

It should be noted, however, that the current study tested and showed that high-value remembered Go items were chosen over high-value forgotten NoGo items, and not that remembered high-value items were chosen over forgotten high-value items (i.e., remembered high-value NoGo items were not always chosen over forgotten high-value Go items; see Figure 4 and Supplementary Figure 5). This finding indicate that memory is not the only process underlying the CAT effect. It is also important to note that our findings are correlational and do not establish a causal effect of memory on choices following CAT. This could have been further tested with a mediation analysis, however unfortunately our design is not suitable for this analysis. Future studies can be designed to test the causal effect of memory on choices in a more direct way.

Our suggested mechanism and our findings, emphasizing the involvement of memory processes in choices, are in line with a few previous theories, such as the preferences as memories framework (PAM) (Weber & Johnson, 2006) and the query theory (Johnson, Häubl, & Keinan, 2007). According to the PAM framework, preferences are the product of memory processes, such that relevant knowledge (rather than some kind of stored “value”) is retrieved from memory when choice alternatives are compared, and thus choices are highly affected by basic memory processes such as priming and reactivation (Weber & Johnson, 2006). Some of the known characteristics of preferences, for instance instability and semantic framing-effects, which might be surprising in the light of other frameworks, are expected when conceptualizing preferences as memory product and representation. The query theory (Johnson et al., 2007) also suggests that preferences are constructed at the time of choice, rather than stored and then retrieved during choices. Query theory further suggests that choices are made based on answers to questions, or queries, which are internally raised sequentially. According to this theory, the order of these queries, which is dependent on the context at the time of choice, substantially affects choices (Johnson et al., 2007). In the context of CAT, better remembered Go items may drive the order of internal queries, such that queries in favor of these items arise first and therefore drive the choice process towards Go items.

Another line of research related to our findings is “memory bias” that have been found in decisions from memory, such that people prefer remembered over forgotten options, even if remembered options are worse than average (unless remembered options are very unattractive) (Gluth, Sommer, Rieskamp, & Buchel, 2015). It has been shown that this memory bias is the result of people’s beliefs that their memory is stronger for better compared to worse options (i.e., that a good option is remembered because it is good, while memory of a bad option is not related to its attractiveness) (Mechera-Ostrovsky & Gluth, 2018). These findings may explain the results of the current study, such that better remembered high-value items are chosen more because participants attribute the better memory of these items to their value, while the memory for better-remembered low-value items is not attributed to value (or even attributed to their unattractiveness, and thus they are not chosen). However, these studies used tasks in which items were paired with specific locations on the screen, and then during the time of choice, these specific locations were presented without the items. Hence, the choice task in those studies forced participants to retrieve the options from memory, and a forgotten option was unknown during choices. This is in contrast to the binary choice task used in CAT, where both items are presented at the time of choice, and the memory measure is related to remembering that a specific item appeared during training (and putatively to its accessibility in memory).

Our hypothesized mechanism suggests that items with greater memory accessibility are chosen more. One possible mechanism for this process was recently proposed (Bakkour et al., 2019; Shadlen & Shohamy, 2016) based on the observation that harder decisions (e.g. between options with similar subjective values) take longer (Krajbich, Armel, & Rangel, 2010; Milosavljevic, Malmaud, Huth, Koch, & Rangel, 2010). In perceptual decision-making, perceptual evidence accumulates with time and a decision is made once a given threshold of evidence has been reached (Gold & Shadlen, 2007; Shadlen & Kiani, 2013). However, during value-based decision-making, it is unclear what consumes the time to decision. Shadlen and Shohamy (Shadlen & Shohamy, 2016) proposed that these decisions are based on retrieval of accumulated evidence from episodic memory. The findings of our current study indicate that enhanced memory accessibility is related to choices, even in the absence of external reinforcements. In the framework of evidence accumulation from episodic memory during value-based decision-making, the relationships we found between choices and memory accessibility may be driven by a faster rate of evidence accumulation from memory in favor of the better remembered Go items.

In conclusion, our findings provide evidence in support of our suggested mechanism underlying the CAT effect, by showing that memory is enhanced for trained items and that enhanced memory is related to subsequent choices. Moreover, they suggest that memory accessibility is involved in value-based choices, even in the absence of external reinforcements. These findings are important for revealing the role of memory in value-based decision-making in general, and specifically in the absence of external reinforcements. These results shed light on the mechanism underlying behavioral change following CAT, which offers novel avenues for long-lasting behavioral change interventions for maladaptive behaviors such as eating disorders and addictions.

## Materials and Methods

### Data Sharing and Pre-Registration

All of our data and codes are publicly shared on GitHub (https://doi.org/10.5281/zenodo.4593642, Release 2.0.0). Pre-registrations can be found on the Open Science Framework (OSF: Experiment 1: https://doi.org/10.17605/OSF.IO/V79MS; Experiment 2: https://doi.org/10.17605/OSF.IO/YH749).

### Participants

A total of *N* = 95 participants took part in this study, each completed one of the three experiments - Experiment 1 (*n* = 35), Experiment 2 (*n* = 35), or a pilot experiment (*n* = 25). See Table 1 for a demographic description of each experimental sample, and the Supplementary Materials for a detailed description of the pilot experiment. We pre-registered the sample sizes of Experiment 1 and Experiment 2. A power analysis performed with our previous CAT samples, yielded a minimal *n* = 20 for detecting the choice effect with 80% power and alpha = .05. However, since the current experiments focused on memory, which was not similarly tested before with CAT, and also taking into account that some participants would probably not return to the follow-up session, we chose to pre-register a larger sample size of *n* = 35 for each experiment.

All participants had normal or corrected-to-normal vision and hearing and provided signed informed consent to participate in the experiment in return for monetary compensation. Participants were asked to refrain from eating and drinking anything but water for one hour prior to each visit in the laboratory. The study was approved by the ethics committee of Tel Aviv University.

### Stimuli

Stimuli in this study comprised of colored images of familiar local Israeli snack food items. Images depicted the snack package and the snack itself on a homogenous black rectangle sized 576 x 432 pixels. The stimuli were created in our laboratory and are available online (http://schonberglab.tau.ac.il/resources/snack-food-image-database/). To promote incentive compatible behavior, when participants entered the laboratory, they were presented with a cabinet containing the real snacks and the items were available for actual consumption at the end of the experiment.

### Procedures

The experiment was run on a 21.5-inch iMAC computer with Matlab version 2014b (Mathworks, Inc. Natick, MA, USA), the Psychtoolbox (http://www.psychtoolbox.org/) and Python-based Pygame package (Python version 2.7).

The general design included an auction (used to obtain subjective willingness to pay), training, a recognition task and a binary choice task (see Figure 1). The tasks, besides the recognition task, were similar to previous studies with CAT (Salomon et al., 2018; Schonberg et al., 2014). The variants of the design for each experiment are described below.

### Subjective preferences evaluation(see Figure 1a)

First, subjective preferences for the snack food items (60 items in Experiment 1 and the Pilot Experiment, and 80 items in Experiment 2) were evaluated using a Becker-DeGroot-Marschak (BDM) auction procedure (Becker et al., 1964). Participants were first endowed with 10 Israeli Shekels (ILS; equivalent to ∼2.7$ US). During the auction, snack food items were individually presented on the screen in a random order. The task was self-paced. Participants were asked to indicate how much money they are willing to pay for each item, using the mouse cursor along a continuous visual analog scale, ranging from 0 - 10 ILS. Participants were informed in advance that at the end of the experiment, the computer will randomly choose one of the items and will generate a counter bid. In case the bid placed by the participant exceeds the computer’s bid, she or he will be required to buy the item for the computer’s lower bid price. Otherwise, the participant will not buy the item and will keep the allocated 10 ILS. Instructions explicitly mentioned that the best strategy for this task was to indicate the actual willingness to pay (WTP) for each item.

### Item selection

For each participant, items were rank ordered based on the subjective WTP values. Items were split to high-value (above the median) and low-value (below the median) items. One set of items within each value category was used as Go items, while another set of items, with identical mean ranks, was used as NoGo items (see Supplementary Table 1). These sets of items were counter-balanced across participants. This item selection procedure allowed us to present choices of high-value Go versus high-value NoGo items, as well as choices of low-value Go versus low-value NoGo items, with similar initial WTPs, in the probe binary phase. Of the items presented during training, 30% were Go items (12 out of the 40 items in Experiment 1 and the Pilot Experiment; 24 out of the 80 items in Experiment 2).

### Training (see Figure 1b)

During the training session, individual items were presented on the screen one by one for one second each. We instructed participants to press the ‘b’ button on the keyboard as fast as they could whenever an auditory Go-cue was heard. Unbeknownst to participants, the cue was consistently paired with Go items and not with NoGo items. The neutral auditory cue, consisted of a 180 ms-long sinus wave function, was initially heard 750 ms after the stimulus onset. Aiming for a success rate of ∼75% (successful button press before the image disappeared from the screen), we updated the delay between the stimulus onset and cue onset with a ladder technique, such that the delay was increased by 16.67 ms following a successful trial and decreased by 50 ms following a missed trial. We used a jittered inter-stimulus interval (ISI) from a truncated exponential distribution, ranging from one to six seconds (one second interval) and an average duration of two seconds. Importantly, no external reinforcement was provided to the participants with regard to their performance on the task.

### Recognition (see Figure 1c)

The recognition task was added to the paradigm in order to test our proposed mechanism. The recognition task was performed either ∼30 minutes after training (Experiment 2) or three days and again one month following training (Experiment 1). In this task, all items that would be included in the subsequent Go-NoGo binary choice probe pairs, as well as an equal number of new items, were presented on the screen one at a time. Participants responded for two consecutive questions: (1) Was the item presented during the experiment and (2) was the item paired with a cue during training. Participants responded using a five-point confidence scale (1 - sure yes, 2 - think yes, 3 - uncertain, 4 - think no, 5 - sure no) within a maximal time frame of three seconds.

### Probe (see Figure 1d)

In the probe phase, we tested participants’ preferences following CAT, using a binary choice task. On each trial, participants were presented with a pair of items from the same value category (both either high-value or low-value), one Go item (i.e. consistently paired with the cue and button-press during training) and one NoGo item (i.e. not paired with the cue and button press during training). On each trial, participants were given 1.5 seconds to choose the item they preferred, by pressing the button ‘u’ (to choose the left item) or the button ‘i’ (to choose the right item) on the keyboard. Participants were told in advance that at the end of the experiment, one trial will be randomly drawn by the computer and that they will receive the item they chose on the randomly drawn trial. The ISIs were sampled from a truncated exponential distribution, ranging from 1 - 12 seconds (one second interval) and an average duration of three seconds.

### Exclusion criteria

Participants were excluded from the final analysis if they placed bids lower than 1 ILS (∼ $0.27 US) on more than 40 items during the BDM auction (Salomon et al., 2018; Schonberg et al., 2014); reached a ladder lower than 200 ms at least once during training, pressed the button without a cue on more than 5% of NoGo training trials (more than 5% false alarm rate) and/or did not press the button following a cue on more than 10% of Go training trials (more than 10% misses). These exclusion criteria were listed in the pre-registration of experiments 1 and 2.

### Analysis

All analyses were performed using R version 3.6.3. For all mixed-effects models, we used R’s glmer and lmer functions (with the packages “lme4” (Bates, Mächler, Bolker, & Walker, 2015) and “lmerTest” (Kuznetsova, 2017)). In all models, random effects were modeled within participants (i.e., we included a random intercept and random slope term per participant). In each model we started with a maximal random effects structure, modeling all random effects and their correlations (Barr, Levy, Scheepers, & Tily, 2013). In case the model did not converge properly, we simplified the maximal model by first removing the random correlations and then reducing the random terms which indicated model converges issues (i.e., correlations of 1, or random variance of 0). Using these criteria preserves type I error while potentially increasing power when random effects estimates are near the boundary values (Matuschek, Kliegl, Vasishth, Baayen, & Bates, 2017).

### Recognition

We predicted that Go stimuli will be better remembered compared to NoGo items (regardless of value category), reflected in higher accuracy (hit rate) and faster response times (RTs) in the old / new recognition task. We only included items that were presented during the probe task in pairs comparing high-value Go versus high-value NoGo or low-value Go versus low-value NoGo items, to ensure that both the number and the mean rank of Go and mean rank of NoGo items within each value category are equal.

Responses in the recognition task were transformed to a binary scale (correct / incorrect), with “uncertain” responses counted as incorrect answers. Missed responses were excluded from analysis. Then, we tested the prediction regarding the hit rate with a one-sided mixed-effects logistic regression with the outcome (correct / incorrect) as the dependent variable and the item type (Go / NoGo) and value category (high-value / low-value) as independent variables. The prediction regarding the RTs was tested with a one-sided mixed-effects linear model with the RT as the dependent variable and the item type (Go / NoGo) and value category (high-value / low-value) as independent variables. In the RT analysis, we included only correct responses, as faster RTs for wrong answers do not imply better memory.

To test for the differential effect at the level of memory modification (i.e., whether Go items are better remembered over NoGo items to a greater extent in pairs of high-value compared to low-value items), in a hierarchical regression, we tested the significance of the interaction between item type (Go / NoGo) and value category (high-value / low-value).

### Choices

We predicted that the CAT effect observed in previous studies will be replicated, such that participants will significantly choose high-value Go over high-value NoGo items (Bakkour et al., 2017; Botvinik-Nezer et al., 2019; Salomon et al., 2018; Schonberg et al., 2014; Veling et al., 2017). Similar to previous studies with the cue-approach task, we tested this prediction using a one-sided mixed-effects logistic regression, comparing the odds of choosing Go items against chance level (log-odds = 0; odds ratio = 1), independently for each value category (high-value and low-value). An interesting aspect of CAT, demonstrated in previous studies with snack food items, was that its effect is greater in choices of high-value Go over NoGo items, than in choices of low-value Go over NoGo items (Botvinik-Nezer et al., 2019; Salomon et al., 2018; Schonberg et al., 2014). Therefore, similar to previous studies, we further tested whether there was a differential effect of higher proportions of Go items choices on high-value compared to low-value probe pairs, with a one-sided mixed-effects logistic regression.

### Relationships between memory and choices

We hypothesized that Go items are preferred over NoGo items because they are more accessible in memory, and therefore their associations accumulate faster to choices and they are chosen more when their associations are positive. Therefore, we predicted that binary choices will be related to memory for the specific alternatives on a trial-by-trial basis, such that better memory for the Go compared to the NoGo item will be related to more Go item choices, mainly for high-value items.

We tested this prediction with two mixed-effects logistic models predicting choices of Go items based on the old / new recognition results: (1) The accuracy model combined the memory for the Go item and the memory for the NoGo item to an independent variable with three accuracy categories: “Go remembered and NoGo forgotten”, “Both remembered / forgotten” and “Go forgotten and NoGo remembered”; (2) The RT model included one independent variable, the *ΔRT* (RT NoGo minus RT Go; only for probe trials in which both items were correctly remembered).

According to our suggested mechanism, better remembered items should be preferred over worse remembered items when associated memories are positive. Therefore, we predicted that this relationship will be stronger for high-value compared to low-value items. We tested this prediction with a mixed-effects logistic regression model with the main effect of value category (high-value / low-value) and the interaction between the value category and the recognition variables. Participants were modelled as a random effect in all models.

### Exploratory analyses

In addition to our pre-registered analyses, we performed the following exploratory analyses (see Supplementary Materials): (1) Comparing RT between Go and NoGo items for incorrect responses (Misses), in order to validate our RT memory effects; (2) comparing hit rate and RTs between Go and NoGo items in the recognition Go / NoGo task; and (3) comparing the confidence levels in the old / new recognition task between Go and NoGo items.

### Deviation from pre-registration

It should be noted that the statistical tests used were correctly pre-registered for Experiment 2 but not for Experiment 1, where we pre-registered we will use t-tests and correlation tests in the recognition analysis and in the analysis of relationships between memory and choices, respectively. However, prior to data analyses, we decided to use the more appropriate mixed-effects tests that are described above (and were pre-registered for Experiment 2, during data collection of Experiment 1). Nonetheless, paired t-tests and linear correlations results are reported in the Supplementary Materials (see section Pre-Registered Analyses for Experiment 1). We also note that our pre-registered analysis plan was not always detailed enough, and some of the models were not explicitly described in enough detail. For example, although our pre-hypothesized suggested mechanism predicts that better remembered items are chosen more when their associations are positive, we did not explicitly include the predictions of the differences between high-value and low-value items in our preregistered analysis plan. These predictions were made prior to analysis and were unintentionally their full detailed were not described in the pre-registration. Furthermore, in Experiment 2 we analyzed the results based on four value categories (high / medium-high / medium-low / low), although this split was not mentioned in our pre-registration, but was used in the original CAT studies (Schonberg et al., 2014). In addition, the pre-registered plan included eye-tracking analysis. While eye-tracking data were indeed collected for some of the participants for exploratory purposes, these data are beyond the scope of this paper, were not analyzed in this study and are not presented here.

### Experiment 1

In Experiment 1, we added a recognition task three days following CAT (i.e., in Session 2). The recognition task was added three days and not immediately following CAT to avoid a ceiling effect that was present in previous experiments in our lab when the recognition task was performed immediately following the probe task (Botvinik-Nezer et al., 2019; Salomon et al., 2018).

Thirty-five valid participants completed the first two sessions, of them *n* = 30 returned for an additional follow-up session, about one month after Session 1 (mean = 34.3 days, *SD* = 5.04 days, range = 28 - 46 days; demographic statistics are reported in Table 1). Four additional participants were excluded based on pre-registered exclusion criteria: two bid less than one ILS on more than 40 items (BDM exclusion criteria) and two reached a ladder lower than 200 ms during training and also pressed the button when not needed (false alarms) on more than 10% of training trials. One participant completed only one of the two probe blocks (i.e., one instead of two repetitions of each probe pair). This participant was included in the analysis; however, the results do not change when this participant is excluded from the analysis.

#### Session 1

Upon arrival, we evaluated participants’ initial preferences for 60 snack food items with a BDM auction (Figure 1a). Then, they completed 16 repetitions of CAT. In each training repetition, 40 items were presented in a random order (Figure 1b).

#### Session 2

Exactly three days after Session 1, participants returned to the laboratory for Session 2. They completed the recognition task (Figure 1c), probe task (Figure 1d) and an additional BDM auction (Figure 1a).

The recognition task included 56 items: 28 new items that were not previously presented during the experiment and 28 old items that were previously presented during the experiment (these would later be used in the probe phase - 24 stimuli in the Go versus NoGo pairs and four additional items from the “sanity check” probe pairs).

Following the recognition task, modifications of preferences were evaluated in a probe task. The probe task consisted of a set of binary choices, in which each of the six high-value Go items was pitted against each of the six high-value NoGo items (6 × 6 = 36 comparisons) and each of the six low-value Go items was pitted against each of the six low-value NoGo items (6 × 6 = 36 comparisons). Thus, overall, there were 72 unique pairs of Go versus NoGo item comparisons. We also included “sanity check” probe pairs, in which high-value NoGo items were pitted against low-value NoGo items. Each unique comparison was repeated twice during the experiment (once in each of two task blocks), resulting in overall 144 Go-NoGo probe trials.

#### Follow-up Session

About one month after Session 1, participants returned to the lab and completed the Follow-up Session, which included the same tasks as in Session 2.

### Experiment 2

Experiment 2 included *n* = 35 valid participants. Three additional participants were excluded based on the pre-registered BDM exclusion criteria. Upon arrival, participants completed a BDM auction with 80 items (instead of 60 items as in Experiment 1), Then, participants completed the training task, which was similar to Experiment 1, besides two important modifications: First, each item was only presented once (instead of 16 repetitions in Experiment 1). Second, 80 (instead of 40) items were presented during training. Out of the 80 items, 12 items were high-value Go items and 12 items were low-value Go items (compared to six high-value Go and six low-value Go items in Experiment 1).

After training, participants completed a filler task during which they ranked fractal art images, unfamiliar faces and familiar faces (of local politicians), as well as the familiarity of the familiar faces. The filler tasks lasted about 30 minutes. Following the filler tasks, participants completed a recognition task, similar to the one performed in Experiment 1, but with a larger total number of 160 items - 80 old and 80 new items.

Then, participants completed the probe task. Since we doubled the number of trained items, we also doubled the number of probe comparisons. Probe Go-NoGo choices included four value categories: high, medium-high, medium-low and low value (for the exact ranks of each value category see Supplementary Table 1). Each category included 36 unique choices between the six Go items pitted against six NoGo items of identical mean rank. Thus, overall, each probe block included a total of 144 unique Go versus NoGo choices. Twelve “sanity check” probe trials (i.e., choices between high-value and low-value NoGo items) were also presented, as in Experiment 1.

Finally, participants completed a familiarity task for the items, in which they ranked the familiarity of each snack food item. However, the results of this task are beyond the scope of this study.

## Acknowledgments

We thank Dr. Jeanette Mumford for her invaluable statistical advice. We thank Alex Manevich, Neta Cohen and Michal Teitler for helping with data collection.

