## Supplementary Materials for "Memory for Individual Items is Related to Non-Reinforced Preference Change"

### Pilot Experiment

In addition to the two pre-registered full experiments described in the main paper, we collected an additional pilot sample to test whether memory is enhanced for Go compared to NoGo items following CAT and whether memory modifications are related to choices. This experiment was similar to Experiment 1, and also consisted of three sessions, but with one important difference: Participants also completed an additional probe task at the end of Session 1.

#### Methods

A total of $n$ = 25 participants completed the first two sessions, $n$ = 14 of them completed the Follow-up Session. Two additional participants were excluded from analysis based on the BDM exclusion criteria (see Exclusion criteria section in general methods).

**Session 1.** Participants completed a BDM auction and a training session, identical to Experiment 1. Following training, participants ranked fractal art images on an analogue scale, as a filler task that lasted about five minutes. Then, we tested their preferences following training with a probe task, which was identical to the probe tasks of Session 2 and the Follow-up Session of Experiment 1.

**Session 2 and Follow-up Session.** These sessions were identical to Session 2 and the Follow-up Session of Experiment 1. Participants returned to Session 2 about four days after Session 1 (interval between days: $M$ = 3.74 days, $SD$ = 1.56 days, *range* = 1 – 7 days) and to the Follow-up Session about one month and a half after Session 1 ($M$ = 45.7 days, $SD$ = 12.6 days, *range* = 27 - 68 days).

#### Results

**Recognition.** ***Session 2.*** (for full statistics see Supplementary Table 10). *Overall performance.* The mean hit rate of participants was 94.75% ($SD$ = 6.42%). The mean correct rejection rate was 79.04% ($SD$ = 13.99%). The mean d' was 2.704 ($SD$ = 0.871). The mean RT was 1.343 (*SD* = 0.195) seconds for hits, 1.816 (*SD* = 0.473) seconds for misses, 1.554 (*SD* = 0.361) seconds for correct rejections and 1.742 (*SD* = 0.356) seconds for false alarms.

*Hit rate.* Hit rate in the old / new recognition task was not significantly higher for Go (*M* = 97.33%) compared to NoGo (*M* = 95.27%) items (*one-sided p* = .095, *odds ratio* = 1.830, 95% *CI* [0.741, 4.515], mixed-effects logistic regression; see Supplementary Figure 3a). Hit rate was also not significantly higher for high-value (*M* = 97.94%) compared to low-value (*M* = 94.60%) items (*one-sided p* = .131, *odds ratio* = 3.103, 95% *CI* [0.430, 22.409]). There was no interaction between the value category (high-value / low-value) and item type (Go / NoGo; *two-sided p* = .897).

*Response time.* RTs in the recognition task were faster for Go (*M* = 1.297 seconds) compared to NoGo (*M* = 1.348 seconds) items, but only with marginal significance (*one-sided p* = .069, *estimated mean difference* = -0.056, mixed-effects linear regression; see Supplementary Figure 3b). RTs were significantly faster for high-value (*M* = 1.287 seconds) compared to low-value (*M* = 1.358 seconds) items (*one-sided p* = .048, *estimated mean difference* = -0.074). There was no interaction between the value category and item type (*two-sided p* = .681).

***Follow-up Session.*** (for full statistics see Supplementary Table 10). *Overall performance.* The mean hit rate of participants was 86.40% (*SD* = 13.43%). The mean correct rejection rate was 54.87% (*SD* = 24.83%). The mean d' was 1.354 (*SD* = 0.757). The mean RT was 1.445 (*SD* = 0.230) seconds for hits, 1.666 (*SD* = 0.414) seconds for misses, 1.558 (*SD* = 0.352) seconds for correct rejections and 1.798 (*SD* = 0.227) seconds for false alarms.

*Hit rate.* Similar to Session 2, hit rate in the old / new recognition task was not significantly higher for Go (*M* = 86.20%) compared to NoGo (*M* = 85.61%) items (*one-sided p* = .458, *odds ratio* = 1.045, 95% *CI* [0.462, 2.364], mixed-effects logistic regression; see Supplementary Figure 3c). Hit rate was also not significantly higher for high-value (*M* =

90.48%) compared to low-value (*M* = 81.21%) items (*one-sided p* = .139, *odds ratio* = 1.789, 95% *CI* [0.624, 5.125]). There was no interaction between the value category and item type (*two-sided p* = .616).

*Response times*. RTs were not significantly faster for Go (*M* = 1.398 seconds) compared to NoGo (*M* = 1.478 seconds) items (*one-sided p* = .085, *estimated mean difference* = -0.070, mixed-effects linear regression; see Supplementary Figure 3d). RTs were not faster for high-value (*M* = 1.464 seconds) compared to low-value (*M* = 1.428 seconds) items (*one-sided p* = .164, *estimated mean difference* = 0.050). There was a significant interaction between the value category and item type, such that RTs were significantly faster for high-value Go compared to high-value NoGo items, but not significantly slower for low-value Go compared to low-value NoGo items (*two-sided p* = .013).

**Choices - all sessions** (for full statistics see Supplementary Table 3). Despite its high replicability in dozens of other samples, the cue-approach effect on preferences was not replicated in this pilot experiment (see Supplementary Figure 4). Participants did not significantly choose high-value Go over high-value NoGo items in all sessions (Session 1: *M* = 52.77%, *one-sided p* = .224; Session 2: *M* = 51.27%, *one-sided p* = .331; Follow-up Session: *M* = 52.96%, *one-sided p* = .216; mixed-effects logistic regression). The proportion of choices of low-value Go over low-value NoGo items was significantly higher than chance level (50%, log-odds=0; odds ratio=1) only in Session 2 (Session 1: *M* = 55.91%, *one-sided p* = .090; Session 2: *M* = 57.07%, *one-sided p* = .031; Follow-up Session: *M* = 47.97%, *one-sided p* = .696). The differential effect of higher proportion of Go items choices on high-value compared to low-value trials was also not significant for all sessions.

**Relationships between memory and choices**. ***Session 2.*** (see Supplementary Table 11). Participants chose Go items significantly more when the Go item was remembered and the NoGo item was forgotten, compared to when both were remembered or forgotten (see

Supplementary Figure 5a), when choosing between high-value items (*one-sided* $p < .001$, *odds ratio* = 4.905, 95% *CI* [1.980, 12.151], mixed-effects logistic regression). The same relationship between recognition accuracy and choices was negative for choices between low-value items (i.e., low-value Go items were chosen less than low-value NoGo items when the Go item was remembered and the NoGo item was forgotten, compared to when both were remembered or forgotten; *one-sided* $p = .973$, *odds ratio* = 0.284, 95% *CI* [0.079, 1.017]). The interaction between this accuracy category and the value category was significant (*one-sided* $p < .001$). There were no significant differences in Go item choices when comparing pairs in which the Go item was forgotten and the NoGo item was remembered, to pairs in which both items were remembered or forgotten, neither for choices between high-value items (*one-sided* $p = .425$, mixed-effects logistic regression) nor for choices between low-value items (*one-sided* $p = .490$). The interaction effect was also not significant (*one-sided* $p = .477$).

The relationship between the ΔRT and choices of high-value Go over high-value NoGo items (see Supplementary Figure 6a) was significantly positive (*one-sided* $p = .002$, *odds ratio* = 2.009, 95% *CI* [1.251, 3.227], mixed-effects logistic regression). This relationship was also significantly positive for low-value probe pairs (*one-sided* $p = .024$, *odds ratio* = 1.557, 95% *CI* [1.004, 2.414]). Although this relationship was stronger for high-value items, the interaction between the ΔRT and value category was not significant (*one-sided* $p = .140$).

***Follow-up Session.*** (see Supplementary Table 12). Participants did not significantly choose Go items more when the Go item was remembered and the NoGo item was forgotten, compared to when both were remembered or forgotten (see Supplementary Figure 5b), neither when choosing between high-value items (*one-sided* $p = .760$, mixed-effects logistic regression) nor when choosing between low-value items (*one-sided* $p = .479$). The interaction between this accuracy category and the value category was also not significant (*one-sided* $p = .158$). Participants significantly chose Go items less when the go item was forgotten and the NoGo item was remembered, compared to when both were remembered or forgotten, when choosing between high-value items (*one-sided p* = .006, *odds ratio* = 0.499, 95% *CI* [0.290, 0.858], mixed-effects logistic regression), but not when choosing between low-value item (*one-sided p* = .092). The interaction was not significant (*one-sided p* = .188).

The relationship between the ΔRT and choices (see Supplementary Figure 6b) of high-value Go over high-value NoGo items was significantly positive (*one-sided p* = .029, *odds ratio* = 2.354, 95% *CI* [0.970, 5.708], mixed-effects logistic regression). This relationship was not significant for low-value probe pairs (*one-sided p* = .120). The interaction between the ΔRT and value category was not significant (*one-sided p* = .166).

### Discussion

In contrast to our predictions and to dozens of previous samples with CAT (Bakkour, Botvinik-Nezer, et al., 2018; Bakkour et al., 2016, 2017; Botvinik-Nezer et al., bioRxiv; Salomon et al., 2018; Schonberg et al., 2014; Veling et al., 2017; Zoltak et al., 2017), in the Pilot Experiment we did not find a significant probe effect of choosing high-value Go over high-value NoGo items. We cannot explain why the CAT effect did not replicate in this specific pilot sample. We did not find differences between the current sample and previous ones, with regard to demographic information, BDM ratings, performance during training or any other aspect of the task.

Analysis of the recognition task revealed that like preferences, memory was also not enhanced for Go compared to NoGo items following CAT in this pilot sample. There are two main possible reasons why memory was not higher for Go compared to NoGo items in recognition task. First, since the overall performance in the recognition task was close to 100%, the ceiling effect may have masked the differences. Second, it is possible that the unknown reason that led to the lack of preference change was also related to the lack of memory change.

Most importantly, we found that memory was related to the specific choices, as in experiments 1 and 2. Furthermore, these relationships were mostly stronger for choices between high-value items compared to choices between low-value items. These results support our hypothesis that for high valued items, with positive associations, better remembered items are chosen over worse remembered items.

### Pre-Registered Analyses for Experiment 1

We originally pre-registered we will test memory differences between Go and NoGo items in Experiment 1 with paired t-tests, and the relationships between memory and choices with linear correlation tests and thus we conducted these analyses. The results are reported below.

#### Testing Recognition Memory With Paired T-Tests

In Session 2, accuracy was not significantly higher for Go compared to NoGo items (paired t-test, *one-sided* $p$ = .096). RTs were significantly faster for Go compared to NoGo items (paired t-test, *one-sided* $p$ = .002). In the Follow-up Session, hit rate was significantly higher (paired t-test, *one-sided* $p$ = .033) and RTs were significantly faster (paired t-test, *one-sided* $p$ = .045) for Go compared to NoGo items.

#### Testing the Relationships Between Memory and Choices With Linear Correlation Tests

In Session 2, the linear correlation between the proportion of choosing high-value Go over high-value NoGo items in the probe task and the difference in hit rate between high-value Go and high-value NoGo items (hit rate for high-value Go minus hit rate for high-value NoGo) in the recognition task, across participants, was not significant (*Pearson's* $r$ = .146, $df$ = 33, *one-sided* $p$ = .201). The linear correlation between the proportion of choosing high-value Go over high-value NoGo items in the probe task and the difference in RTs between high-value NoGo and high-value Go items (RT for high-value NoGo minus RT for high-value Go) in the recognition task, across participants, was also not significant (*Pearson's* $r$ = .204, $df$ = 33, *one-sided* $p$ = .120). In the Follow-up Session, the linear correlation with the hit rate difference across participants was not significant (*Pearson's* $r$ = .027, $df$ = 28, *one-sided* $p$ = .444). The linear correlation with the difference in RTs across participants was significant (*Pearson's* $r$ = .613, $df$ = 28, *one-sided* $p$ < .001).

### Supplementary Exploratory Analyses

#### RT Differences for Incorrect Responses

As predicted, in Experiment 1, we found that RTs in the old / new recognition task were faster for Go compared to NoGo items, when the response was correct (i.e., the participant correctly recognized the item as an old item). We interpreted these results as an indication of stronger memory for Go compared to NoGo items. However, this difference might have been the result of the association between Go items and button pressing established during training, which may lead to faster responses when Go items are presented. In order to test this alternative explanation, we compared the RTs for Go versus NoGo items in incorrect responses during the old / new recognition task. If responses to Go items were faster due to the association between these items and motor responses, independent of memory strength, incorrect responses should also be faster for Go compared to NoGo items. To test this, we took the recognition data from Experiment 1 (where RT differences were found) and compared the difference in RT for Go versus NoGo items, for misses and hits. We also tested the interaction between the item type (Go / NoGo) and the response accuracy (hit / miss), with a mixed-effects linear regression (participants were modelled as a random effect in all our analyses).

Responses were significantly faster for Go compared to NoGo items in correct trials (Go: $M = 1.404$, $SD = 0.277$; NoGo: $M = 1.495$, $SD = 0.262$; *one-sided* $p = .002$), but not in incorrect trials (Go: $M = 1.678$, $SD = 0.565$; NoGo: $M = 1.663$, $SD = 0.487$; *one-sided* $p = .437$). Although descriptively the RT difference was larger for correct versus incorrect responses, the interaction was not significant (*one-sided* $p = .304$). It should be noted that the hit rates in this experiment were high, and thus the number of misses was relatively low. Compared to 695 correct responses (hits) for Go items and 662 hits for NoGo items, there were only 101 incorrect responses (misses) for Go items and 143 misses for NoGo items. These large differences in the number of trials limit our interaction analysis.

**Go / NoGo Recognition Task**

We did not include the results of the Go / NoGo recognition task (the second question shown for each item in the recognition task for all experiments) in our confirmatory analyses. Our hypothesized mechanism does not suggest predictions regarding this task, since although correct recognition of items as Go or NoGo items indicate better memory in general, it does not necessarily relate to better memory accessibility. Moreover, this study was focused on testing memory differences between Go and NoGo items following CAT and their relationships to choices. Comparing hit rate or RTs of Go versus NoGo items in the Go / NoGo recognition is problematic, since the correct responses for each item type, as well as the base rate, are different.

Nevertheless, we describe here the results of this task (See Supplementary Table 13 for full statistics). It should be noted that there were many more NoGo compared to Go items (all new items as well as 16 out of the 28 old items were NoGo items). The following measurements were computed across all items in the recognition task, both new and old. Across all experiments and sessions, the mean hit rate across participants in the Go / NoGo recognition task was relatively low (range 27% - 55%), while the correct rejection rate was higher (range 57.8% - 72.4%). The mean d' was relatively low as well (range -0.2 – 1).

We further tested whether responses were faster for Go compared to NoGo items in the Go / NoGo recognition task, in Experiment 1 (as we did for the old / new recognition task). RTs were significantly faster for Go compared to NoGo items in correct responses (hits / correct rejections. Go: $M = 0.799$, $SD = 0.363$; NoGo: $M = 0.891$, $SD = 0.364$; *two-sided* $p = .008$), and insignificantly slower for Go compared to NoGo items in incorrect responses (misses / false alarms. Go: $M = 0.950$, $SD = 0.310$; NoGo: $M = 0.892$, $SD = 0.326$; *two-sided* $p = .496$). The interaction between item type (Go / NoGo) and correct / incorrect responses was significant (*two-sided* $p = .005$). Since hit rates in this task were not as large as the old / new recognition task, the number of trials in each of these categories were relatively similar (Correct responses: 398 trials with Go items and 387 trials with NoGo items; incorrect responses: 383 trials with Go items and 401 trials with NoGo items).

**Confidence Levels**

Our hypothesized mechanism suggests that Go items are better remembered than NoGo items following CAT. In the recognition task, participants indicated their confidence level on each trial by choosing one of five possible responses (i.e., 1- sure yes, 2- think yes, 3- uncertain, 4- think no, 5- sure no). Therefore, for each answer besides uncertain, the participant had either high or low confidence. Across our confirmatory analyses, we converted these five possible responses to binary responses (correct / incorrect), while ignoring the confidence level. Here, we compared the confidence level for correct responses in the old / new recognition task, between Go and NoGo items (including only the items that were presented in the probe task, as we did for the previous analyses beside the general recognition performance). Across all experiments and sessions, participants were descriptively more certain when recognizing Go items as old items than when recognizing NoGo items as old items (see Supplementary Table 14). However, this difference was only significant in Session 2 of Experiment 1 (*two-sided* $p = .044$). These differences, although exploratory, provide further evidence in favor of stronger memory for Go compared to NoGo items following CAT.

**Supplementary Figures**

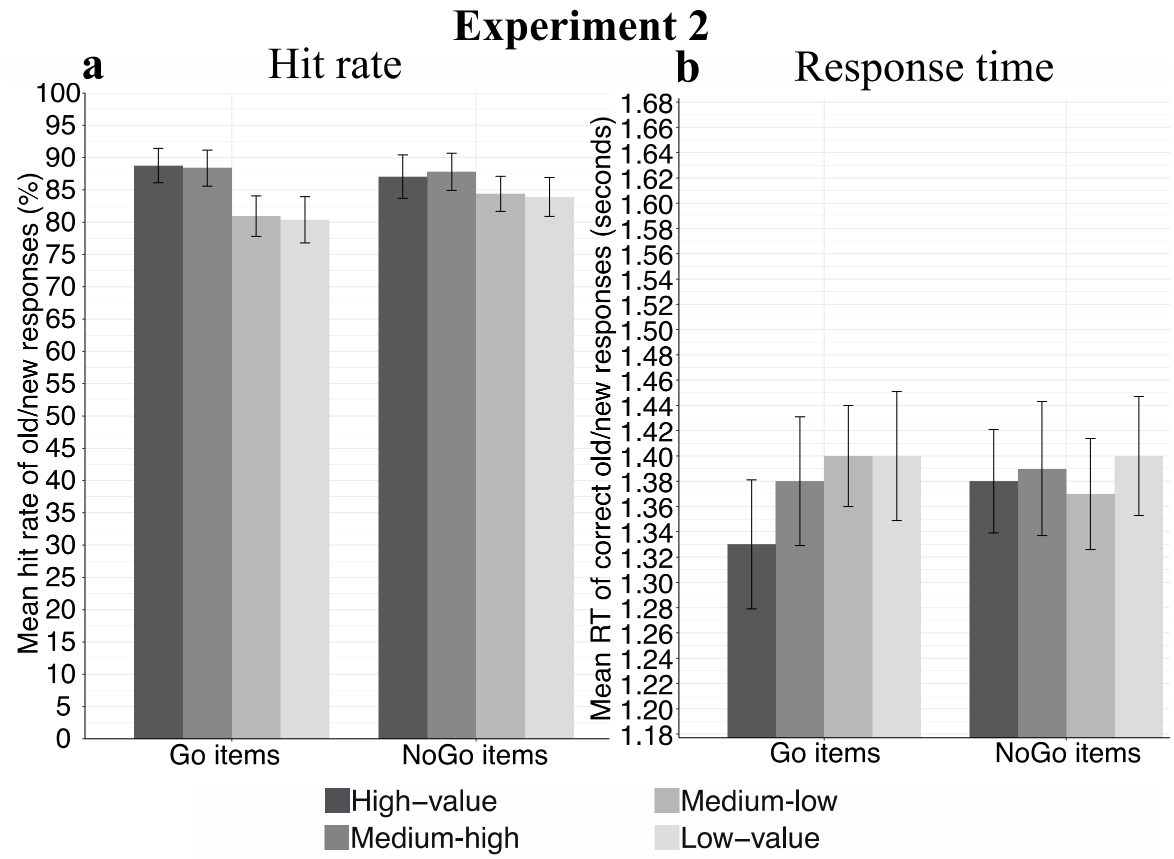


*Supplementary Figure 1*. Recognition result of Experiment 2: (a) Hit rate and (b) RT. The mean of each participant was calculated and then averaged across participants. Error bars represent standard error of the mean across participants.

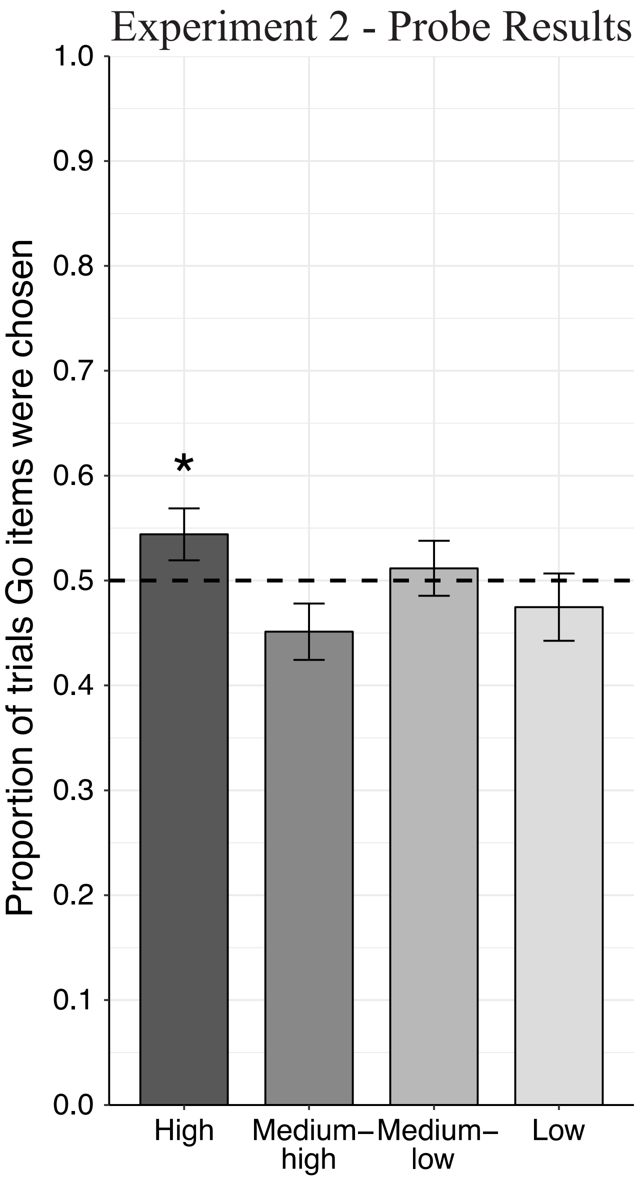


*Supplementary Figure 2*. Probe results of Experiment 2. Mean proportions of choosing the Go item (calculated for each participant and then averaged across participants) are presented with error bars representing standard error of the mean. Dashed line indicates 50% chance level. Asterisks represent statistical significance of a one-sided logistic regression analysis; * $p < .05$

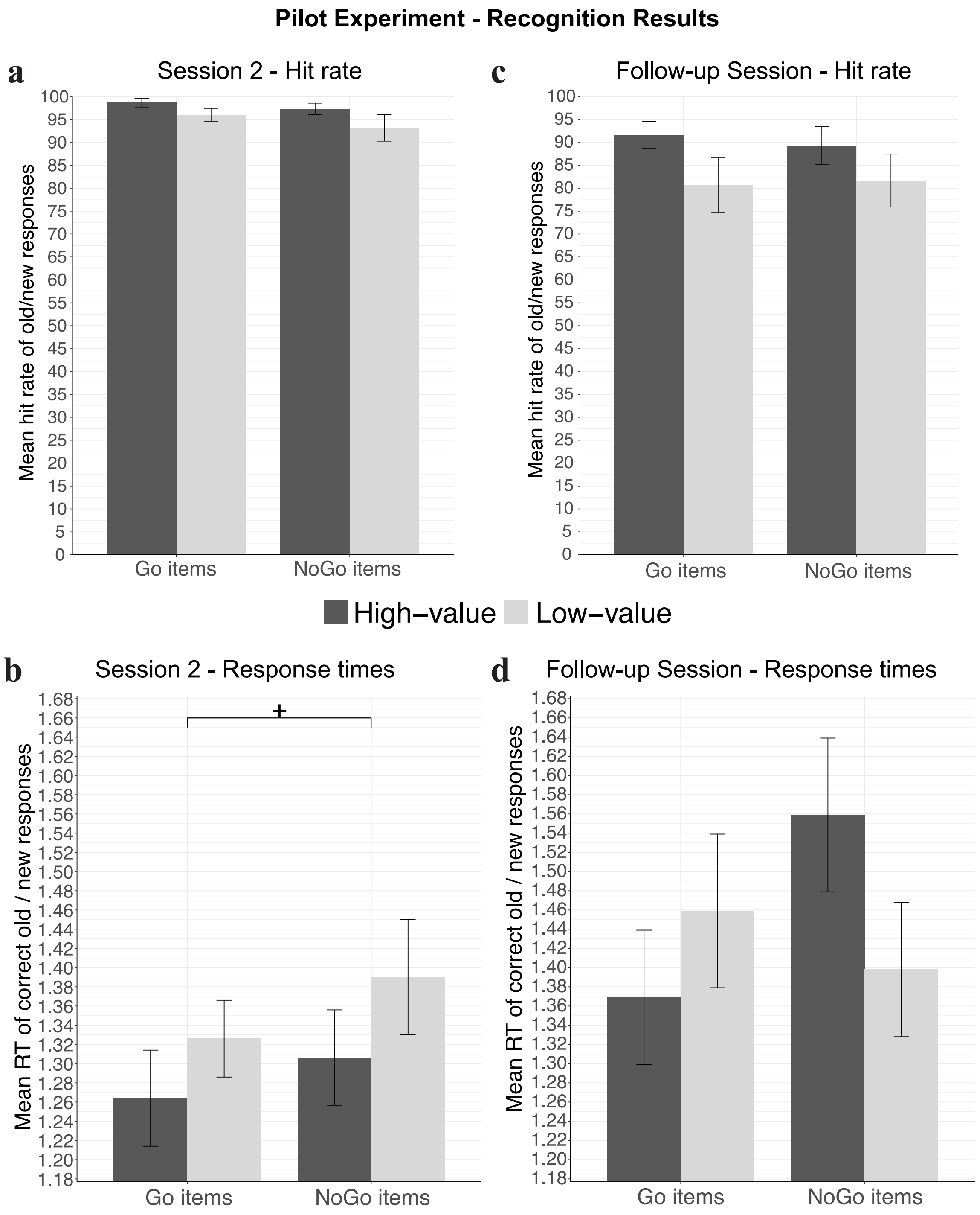


*Supplementary Figure 3.* Recognition results of the Pilot Experiment. The mean hit rate (a,c) and response times (b,d) of each participant were calculated and then averaged across participants. Error bars represent standard error of the mean across participants. Asterix represent one-sided statistical significance; + $p < .07$

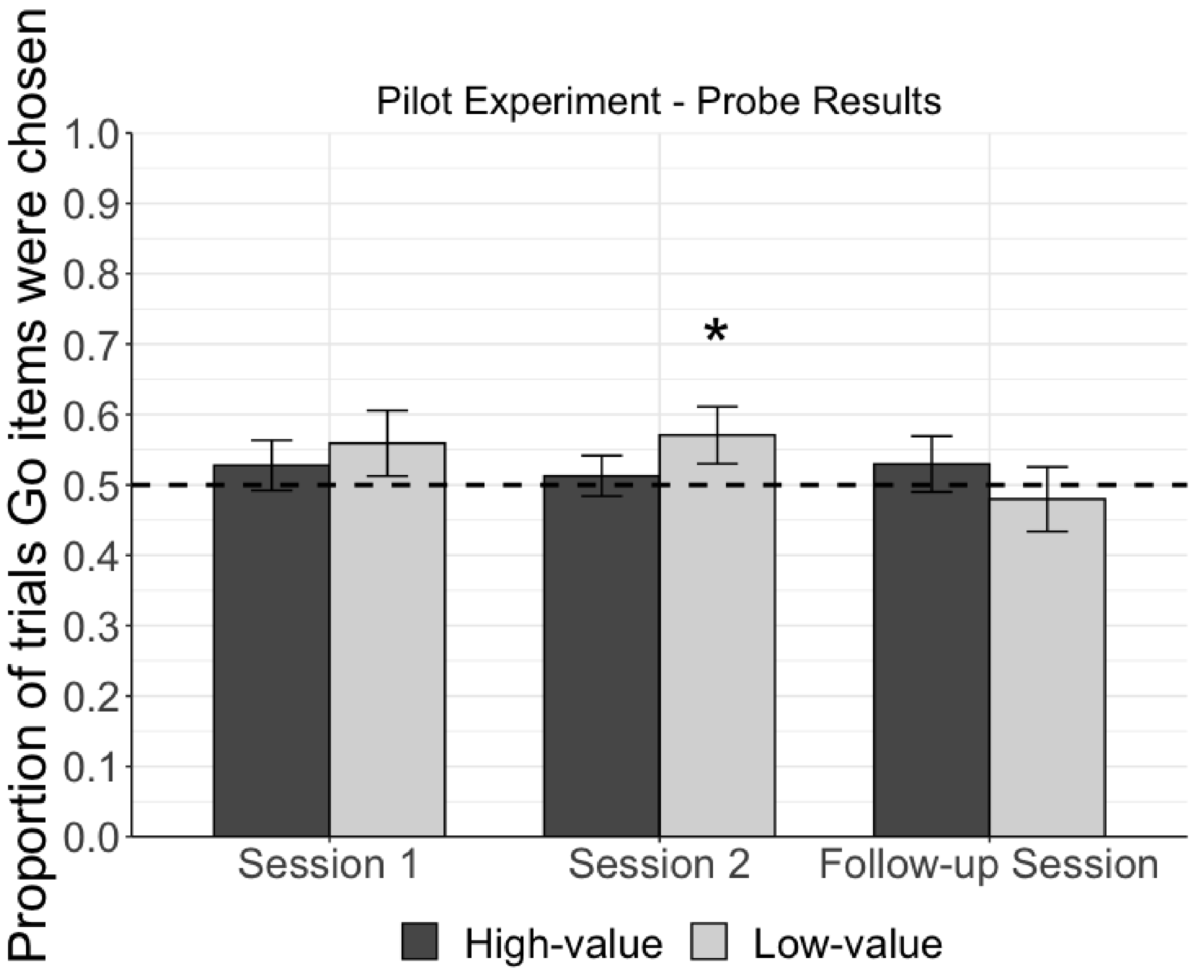


*Supplementary Figure 4*. Probe results of the pilot experiment. Mean proportions of choosing the Go item (calculated for each participant and then averaged across participants) are presented with error bars representing standard error of the mean. Dashed line indicates 50% chance level. Asterisks represent statistical significance of a one-sided logistic regression analysis; * $p < .05$

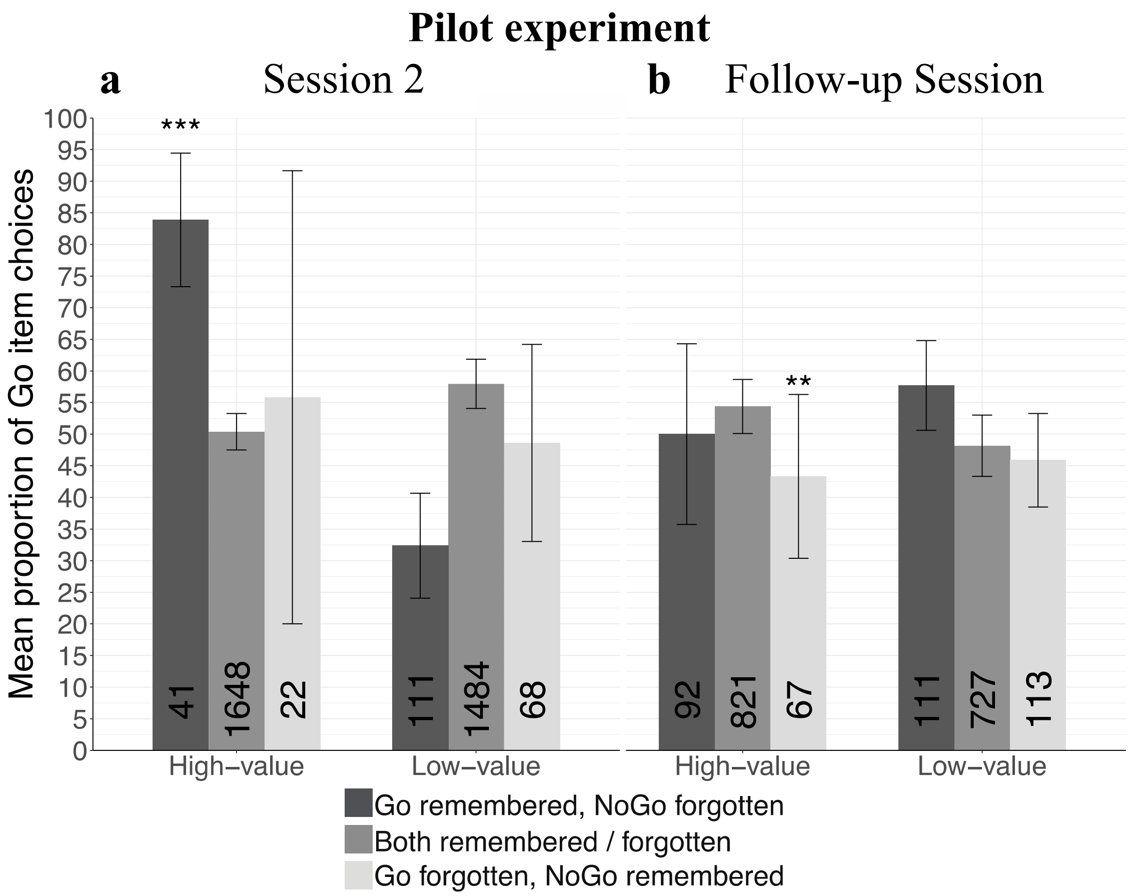


*Supplementary Figure 5*. The relationships between recognition memory and choices in the Pilot Experiment for (a) Session 2 (performed a few days after Session 1) and (b) the Follow-up Session (performed about one month and a half after Session 1). Mean percent of Go item choices as a function of recognition memory with error bars representing standard error of the mean. The number of trials, summed across participants, is presented at the bottom of each bar. Asterisks represent statistical significance between each category and the "Both remembered / forgotten" baseline category (one-sided mixed-effects logistic regression); ** $p < .01$, *** $p < .001$.

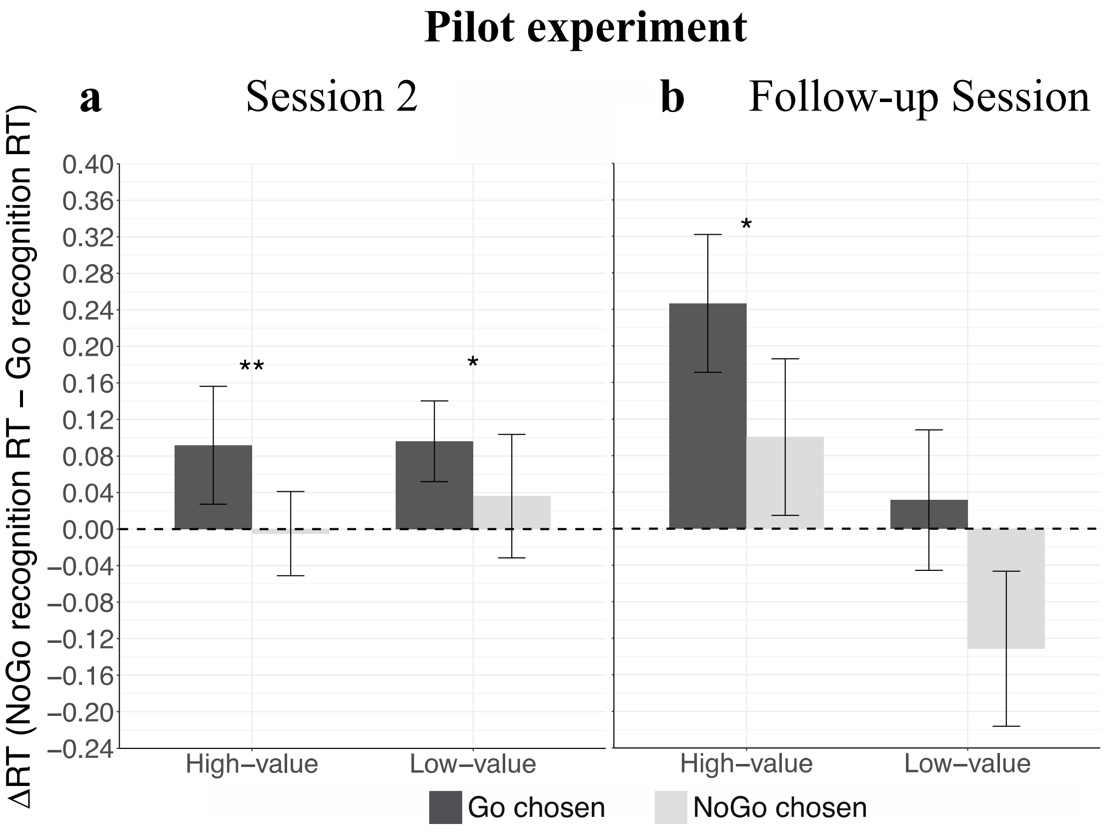


*Supplementary Figure 6. T*he relationships between recognition ΔRT and choices in the pilot experiment, for (a) Session 2 (performed a few days after Session 1) and (b) the Follow-up Session (performed about one month and a half after Session 1). Mean ΔRT values (in seconds) are presented for choices of Go items and for choices of NoGo items, within each value category. Error bars represent standard error of the mean. Asterisks represent statistical significance of one-sided logistic regression; * $p < .05$, ** $p < .01$.

### Supplementary Tables

Supplementary Table 1
*Item Allocation to the Value Categories*

| | **Experiment 1 and Pilot Experiment** | **Experiment 2** |
|---|---|---|
| High-value Go items | 7, 10, 12, 13, 15, 18 | High: 7, 10, 12, 13, 15, 18<br>Medium-high: 24, 25, 27, 30, 32, 33 |
| High-value NoGo items | 8, 9, 11, 14, 16, 17 | High: 8, 9, 11, 14, 16, 17<br>Medium-high: 23, 26, 28, 29, 31, 34 |
| Low-value Go items | 44, 45, 47, 50, 52, 53 | Medium-low: 47, 50, 52, 53, 55, 58<br>Low: 64, 65, 67, 70, 72, 73 |
| Low-value NoGo items | 43, 46, 48, 49, 51, 54 | Medium-low: 48, 49, 51, 54, 56, 57<br>Low: 63, 66, 68, 69, 71, 74 |
| NoGo filler items[a] | 3-6, 19-22, 39-42, 55-58 | 1-6, 19-22, 35-46, 59-62, 75-80 |

*Note.* Rank order of the items are shown for each experiment and set of items (rank 1 = highest). In each experiment, item selection was counter-balanced across participants, such that for half of the participants the items were divided to groups as described in this table, while for the other half the Go and NoGo sets of items were switched (e.g. in Experiment 1, for the other half high-value Go items were items 8, 9, 11, 14, 16, 17 while high-value NoGo items were items 7,10, 12, 13, 15, 18). [a]NoGo filler items were included in training to maintain a proportion of 30% Go items, but were not presented in the Go-NoGo comparisons during probe.

Supplementary Table 2
*Statistics for the Recognition Task of Experiment 1.*

| **Item type** | **Value category** | **Session 2** | | **Follow-up Session** | |
|---|---|---|---|---|---|
| | | **Hit rate** | **RT** | **Hit rate** | **RT** |
| Go | High-value | 87.14% (20.34%) | 1.397 (0.33) | 91.11% (15.62%) | 1.410 (0.36) |
| | Low-value | 90.10% (17.16%) | 1.391 (0.30) | 86.28% (18.66%) | 1.444 (0.33) |
| | All | 88.51% (15.18%) | 1.392 (0.29) | 88.81% (14.06%) | 1.422 (0.33) |
| NoGo | High-value | 85.14% (15.81%) | 1.503 (0.28) | 84.22% (16.00%) | 1.481 (0.36) |
| | Low-value | 84.38% (13.97%) | 1.521 (0.30) | 82.89% (15.41%) | 1.473 (0.34) |
| | All | 84.97% (9.77%) | 1.514 (0.27) | 83.53% (13.99%) | 1.477 (0.34) |
| All | High-value | 86.30% (13.75%) | 1.448 (0.28) | 87.57% (13.58%) | 1.442 (0.34) |
| | Low-value | 87.40% (11.65%) | 1.454 (0.26) | 84.77% (13.10%) | 1.455 (0.32) |
| | All | 86.80% (9.99%) | 1.450 (0.26) | 86.15% (11.73%) | 1.447 (0.32) |

*Note.* Hit rate and mean response time (RT, in seconds) in the old / new recognition task for the Go and NoGo items that were compared during the probe task, were calculated for each participant, and then averaged across participants. Means are presented with standard deviation (across participants) in brackets.

Supplementary Table 3

*Probe Results of Experiment 1 and the pilot experiment*

| **Experiment** | **Session** | **Value category** | **Mean** | **SEM** | ***p*** | **OR** | **95% CI** |
|---|---|---|---|---|---|---|---|
| **Experiment 1** | **Session 2** | High | 61.67% | 2.9% | < **.001** | 1.745 | 1.647 - 1.849 |
| | | Low | 58.16% | 3.6% | **.009** | 1.487 | 1.387 - 1.594 |
| | **Follow-up** | High | 59.62% | 2.8% | < **.001** | 1.546 | 1.464 - 1.632 |
| | | Low | 54.95% | 3.4% | .066 | 1.244 | 1.164 - 1.330 |
| **Pilot Experiment** | **Session 1** | High | 52.77% | 3.6% | .224 | 1.123 | 1.048 - 1.205 |
| | | Low | 55.91% | 4.7% | .090 | 1.332 | 1.216 - 1.459 |
| | **Session 2** | High | 51.27% | 2.9% | .331 | 1.054 | 0.996 - 1.115 |
| | | Low | 57.07% | 4.1% | **.031** | 1.420 | 1.312 - 1.538 |
| | **Follow-up** | High | 52.96% | 4.0% | .216 | 1.136 | 1.051 - 1.228 |
| | | Low | 47.97% | 4.6% | .696 | 0.905 | 0.827 - 0.990 |

*Note.* Statistics for the binary choices (probe) phase. Percent of Go item choices were calculated for each participant, and then averaged across participants. Means are presented with standard errors (SEM; across participants). One-sided p-values (logistic regression) are presented with the odds ratio (OR) and 95% confidence interval (CI).

Supplementary Table 4

*Statistics for the Relationships Between Memory and Choices in Experiment 1, Session 2.*

| **Value category** | **Accuracy model: Go remembered, NoGo forgotten** | | | **Accuracy model: Go forgotten, NoGo remembered** | | | **RT model: ΔRT ($RT_{NoGo}$ - $RT_{Go}$)** | | |
|---|---|---|---|---|---|---|---|---|---|
| | ***p*** | **OR** | **95% CI** | ***p*** | **OR** | **95% CI** | ***p*** | **OR** | **95% CI** |
| High-value | **.026** | 2.258 | 0.993-5.135 | .102 | 0.658 | 0.345-1.255 | .171 | 1.219 | 0.81-1.833 |
| Low-value | **.024** | 1.341 | 1.003-1.792 | .893 | 1.265 | 0.873-1.834 | .302 | 1.133 | 0.706-1.821 |
| Interaction with value category | **.044** | 1.545 | 0.937-2.549 | .088 | 0.676 | 0.383-1.193 | .359 | 1.06 | 0.773-1.453 |

*Note.* One-sided p-values are shown for each independent variable (or category), as well as the odds ratio (OR; which indicate how much larger the odds of choosing the Go items are with the change of category in the accuracy model, or each increase of 1 unit in the RT model) and the 95% confidence interval of the odds ratio (CI). The dependent variable in the logistic regression analysis is Go item choices (1- chose the Go item, 0- chose the NoGo item).

Supplementary Table 5

*Statistics for the Relationships Between Memory and Choices in Experiment 1, Follow-up Session.*

| Value category | Accuracy model: Go remembered, NoGo forgotten | | | Accuracy model: Go forgotten, NoGo remembered | | | RT model: ΔRT ($RT_{NoGo}$ - $RT_{Go}$) | | |
|---|---|---|---|---|---|---|---|---|---|
| | ***p*** | **OR** | **95% CI** | ***p*** | **OR** | **95% CI** | ***p*** | **OR** | **95% CI** |
| High-value | **< .001** | 2.810 | 1.774-4.450 | .589 | 1.059 | 0.642-1.747 | .174 | 1.340 | 0.727-2.471 |
| Low-value | .237 | 1.315 | 0.622-2.780 | .503 | 1.005 | 0.342-2.951 | .924 | 0.726 | 0.469-1.125 |
| Interaction with value category | **.002** | 2.037 | 1.258-3.299 | .454 | 0.955 | 0.433-2.107 | **.005** | 1.551 | 1.110-2.166 |

*Note.* One-sided p-values are shown for each independent variable (or category), as well as the odds ratio (OR; which indicate how much larger the odds of choosing the Go items are with the change of category in the accuracy model, or each increase of 1 unit in the RT model) and the 95% confidence interval of the odds ratio (CI). The dependent variable in the logistic regression analysis is Go item choices (1- chose the Go item, 0- chose the NoGo item).

Supplementary Table 6
*Statistics for the Recognition Task of Experiment 2.*

| Item type | Value category | Hit rate (percent) | RT (seconds) |
|---|---|---|---|
| Go | High | 88.76% (15.76%) | 1.330 (0.299) |
| | Medium-high | 88.38% (16.54%) | 1.384 (0.303) |
| | Medium-low | 80.95% (18.59%) | 1.400 (0.237) |
| | Low | 80.38% (21.19%) | 1.400 (0.300) |
| | All | 84.67% (12.28%) | 1.380 (0.232) |
| NoGo | High | 87.05% (19.87%) | 1.375 (0.241) |
| | Medium-high | 87.81% (17.05%) | 1.387 (0.314) |
| | Medium-low | 84.38% (16.14%) | 1.371 (0.260) |
| | Low | 83.90% (17.79%) | 1.395 (0.279) |
| | All | 85.78% (14.42%) | 1.383 (0.221) |
| All | High | 87.91% (15.97%) | 1.350 (0.225) |
| | Medium-high | 88.16% (14.45%) | 1.386 (0.266) |
| | Medium-low | 82.55% (15.16%) | 1.386 (0.213) |
| | Low | 82.11% (15.66%) | 1.393 (0.251) |
| | All | 85.20% (12.46%) | 1.380 (0.214) |

*Note.* Hit rate and mean response time (RT) in the old / new recognition task for the Go and NoGo items that were compared during the probe task, were calculated for each participant, and then averaged across participants. Means are presented with standard deviations (across participants) in brackets.

Supplementary Table 7

*Probe Results of Experiment 2*

| Value category | Mean | SEM | *p* | OR | 95% CI |
|---|---|---|---|---|---|
| **High** | 54.41% | 2.50% | **.035** | 1.209 | 1.152 - 1.270 |
| **Medium-high** | 45.13% | 2.70% | .963 | 0.814 | 0.773 - 0.858 |
| **Medium-low** | 51.17% | 2.60% | .334 | 1.048 | 0.996 - 1.103 |
| **Low** | 47.47% | 3.20% | .756 | 0.905 | 0.850 - 0.964 |

*Note.* Statistics for the binary choices (probe) phase of Experiment 2. Percent of Go item choices were calculated for each participant, and then averaged across participants. Standard errors (SEM) were computed across participants. One-sided p-values (logistic regression) are presented, as well as the odds ratio (OR) and 95% confidence interval (CI).

Supplementary Table 8

*Statistics for the Relationships Between Memory and Choices in Experiment 2.*

| **Value category** | **Accuracy model** | | | | | | **RT model** | | |
|---|---|---|---|---|---|---|---|---|---|
| | **Go remembered, NoGo forgotten** | | | **Go forgotten, NoGo remembered** | | | **ΔRT ($RT_{NoGo}$ - $RT_{Go}$)** | | |
| | ***p*** | **OR** | **95% CI** | ***p*** | **OR** | **95% CI** | ***p*** | **OR** | **95% CI** |
| High-value | **.004** | 2.627 | 1.275-5.410 | **.009** | 0.499 | 0.280-0.888 | **.004** | 1.511 | 1.108-2.059 |
| The rest | **.016** | 1.287 | 1.023-1.619 | .270 | 0.936 | 0.759-1.156 | **.002** | 1.371 | 1.101-1.708 |
| Interaction with value category (high vs. the rest) | **.016** | 1.727 | 1.047-2.849 | **.009** | 0.527 | 0.310-0.896 | .329 | 1.073 | 0.785-1.469 |

*Note.* One-sided p-values are shown for each independent variable (or category), as well as the odds ratio (OR; which indicate how much larger the odds of choosing the Go items are with the change of category in the accuracy model, or each increase of 1 unit in the RT model) and the 95% confidence interval of the odds ratio (CI). The dependent variable in the logistic regression analysis is Go item choices (1- chose the Go item, 0- chose the NoGo item).

Supplementary Table 9

*Statistics for the Relationships Between Memory and Choices in Experiment 2, for "the Rest" Value Categories*

| | Accuracy model | | | | | | RT model | | |
|---|---|---|---|---|---|---|---|---|---|
| **Value category** | **Go remembered, NoGo forgotten** | | | **Go forgotten, NoGo remembered** | | | **ΔRT ($RT_{NoGo}$ - $RT_{Go}$)** | | |
| | ***p*** | **OR** | **95% CI** | ***p*** | **OR** | **95% CI** | ***p*** | **OR** | **95% CI** |
| Medium-high | .398 | 1.110 | 0.501-2.458 | .381 | 0.913 | 0.505-1.650 | **.036** | 1.370 | 0.972-1.931 |
| Medium-low | .227 | 1.219 | 0.727-2.045 | **.041** | 0.640 | 0.388-1.057 | **.001** | 1.896 | 1.245-2.887 |
| Low | **.047** | 1.438 | 0.941-2.197 | .271 | 1.123 | 0.772-1.634 | **.045** | 1.394 | 0.950-2.046 |

*Note.* One-sided p-values are shown for each independent variable, as well as the odds ratio (OR; which indicate how much larger the odds of choosing the Go items are with each increase of 1 unit in the independent variable) and the 95% confidence interval (CI). The dependent variable in the logistic regression analysis is Go item choices (1- chose the Go item, 0- chose the NoGo item).

Supplementary Table 10

*Statistics for the Recognition Task of the Pilot Experiment.*

| **Item type** | **Value category** | **Session 2** | | **Follow-up Session** | |
|---|---|---|---|---|---|
| | | **Hit rate** | **RT** | **Hit rate** | **RT** |
| Go | High-value | 98.67% (4.61%) | 1.264 (0.24) | 91.67% (10.84%) | 1.369 (0.27) |
| | Low-value | 96.00% (7.26%) | 1.326 (0.19) | 80.71% (22.50%) | 1.459 (0.29) |
| | All | 97.33% (4.64%) | 1.297 (0.20) | 86.20% (13.67%) | 1.398 (0.25) |
| NoGo | High-value | 97.33% (6.24%) | 1.306 (0.24) | 89.29% (15.48%) | 1.559 (0.30) |
| | Low-value | 93.20% (14.54%) | 1.390 (0.28) | 81.67% (21.59%) | 1.398 (0.28) |
| | All | 95.27% (8.79%) | 1.348 (0.22) | 85.61% (14.80%) | 1.478 (0.26) |
| All | High-value | 97.94% (4.45%) | 1.287 (0.21) | 90.48% (11.26%) | 1.464 (0.26) |
| | Low-value | 94.60% (7.97%) | 1.358 (0.20) | 81.21% (20.32%) | 1.428 (0.25) |
| | All | 96.26% (4.93%) | 1.323 (0.19) | 85.91% (13.22%) | 1.438 (0.23) |

*Note.* Hit rate and mean response time (RT, in seconds) in the old / new recognition task for the Go and NoGo items that were compared during the probe task, were calculated for each participant, and then averaged across participants. Means are presented with standard deviation (across participants) in brackets.

Supplementary Table 11

*Statistics for the Relationships Between Memory and Choices in the Pilot Experiment, Session 2.*

| **Value category** | **Accuracy model** | | | | | | **RT model** | | |
|---|---|---|---|---|---|---|---|---|---|
| | **Go remembered, NoGo forgotten** | | | **Go forgotten, NoGo remembered** | | | **ΔRT ($RT_{NoGo}$ - $RT_{Go}$)** | | |
| | ***p*** | **OR** | **95% CI** | ***p*** | **OR** | **95% CI** | ***p*** | **OR** | **95% CI** |
| High-value | >**.001** | 4.905 | 1.980-12.151 | .425 | 0.914 | 0.361-2.314 | **.002** | 2.009 | 1.251-3.227 |
| Low-value | .973 | 0.284 | 0.079-1.017 | .490 | 0.976 | 0.156-6.109 | **.024** | 1.557 | 1.004-2.414 |
| Interaction with value category | >**.001** | 11.253 | 3.994-31.707 | .477 | 1.033 | 0.348-3.064 | .140 | 1.220 | 0.850-1.751 |

*Note.* One-sided p-values are shown for each independent variable (or category), as well as the odds ratio (OR; which indicate how much larger the odds of choosing the Go items are with the change of category in the accuracy model, or each increase of 1 unit in the RT model) and the 95% confidence interval of the odds ratio (CI). The dependent variable in the logistic regression analysis is Go item choices (1- chose the Go item, 0- chose the NoGo item).

Supplementary Table 12

*Statistics for the Relationships Between Memory and Choices in the Pilot Experiment, Follow-up Session.*

| **Value category** | **Accuracy model** | | | | | | **RT model** | | |
|---|---|---|---|---|---|---|---|---|---|
| | **Go remembered, NoGo forgotten** | | | **Go forgotten, NoGo remembered** | | | **ΔRT ($RT_{NoGo}$ - $RT_{Go}$)** | | |
| | ***p*** | **OR** | **95% CI** | ***p*** | **OR** | **95% CI** | ***p*** | **OR** | **95% CI** |
| High-value | .760 | 0.840 | 0.517-1.364 | **.006** | 0.499 | 0.290-0.858 | **.029** | 2.354 | 0.970-5.708 |
| Low-value | .479 | 1.013 | 0.635-1.615 | .092 | 0.735 | 0.467-1.157 | .120 | 1.393 | 0.802-2.421 |
| Interaction with value category | .158 | 1.652 | 0.620-4.401 | .188 | 0.704 | 0.323-1.532 | .166 | 1.286 | 0.774-2.137 |

*Note.* One-sided p-values are shown for each independent variable, as well as the odds ratio (OR; which indicate how much larger the odds of choosing the Go items are with each increase of 1 unit in the independent variable) and the 95% confidence interval of the odds ratio (CI). The dependent variable in the logistic regression analysis was Go item choices (1- chose the Go item, 0- chose the NoGo item).

Supplementary Table 13

*Descriptive Statistics for the Go / NoGo Recognition Task*

| **Experiment** | **Session** | **Hit rate** | **Correct rejection rate** | **d'** |
|---|---|---|---|---|
| Experiment 1 | Session 2 | 54.52% (26.30%) | 72.44% (23.69%) | 1.003 (1.369) |
| | Follow-up | 46.89% (26.33%) | 61.16% (23.01%) | 0.205 (1.426) |
| Experiment 2 | - | 27.01% (12.02%) | 66.06% (11.25%) | -0.249 (0.616) |
| pilot experiment | Session 2 | 54.99% (24.89%) | 67.71% (20.00%) | 0.774 (1.294) |
| | Follow-up | 34.85% (16.76%) | 57.82% (23.89%) | -0.209 (1.234) |

Supplementary Table 14

*Descriptive Statistics for the Confidence Levels in the Old / New Recognition Task*

| **Experiment** | **Session** | **Confidence Go items (1-high; 0-low) Mean (SD)** | **Confidence NoGo items (1-high; 0-low) Mean (SD)** | ***p*** | **OR** | **95 %CI** |
|---|---|---|---|---|---|---|
| Experiment 1 | Session 2 | 0.845 (0.209) | 0.792 (0.210) | **.044** | 2.123 | 1.021-4.418 |
| Experiment 1 | Follow-up | 0.700 (0.290) | 0.645 (0.278) | .089 | 1.419 | 0.948-2.125 |
| Experiment 2 | - | 0.876 (0.158) | 0.871 (0.158) | .781 | 1.056 | 0.720-1.547 |
| Pilot Experiment | Session 2 | 0.954 (0.079) | 0.932 (0.112) | .938 | 1.062 | 0.237-4.762 |
| Pilot Experiment | Follow-up | 0.709 (0.199) | 0.695 (0.194) | .831 | 1.061 | 0.617-1.822 |

*Note.* Statistics are shown for the exploratory confidence levels analysis of each experiment and session. Mean confidence level was computed across participants and is shown with standard deviation (*SD*) values across participants in brackets. Two-sided *p*-values are presented with the odds ratio (*OR*) and the 95% confidence interval of the odds ratio (*CI*). The dependent variable in the logistic regression analysis was whether the confidence level was high or low (1- high, 0- low).
